# Redox potential defines functional states of adult hippocampal stem cells

**DOI:** 10.1101/606186

**Authors:** Vijay S Adusumilli, Tara L Walker, Rupert W Overall, Gesa M Klatt, Salma A Zeidan, Tim J Fischer, Sara Zocher, Alex M Sykes, Susanne Reinhardt, Andreas Dahl, Dilyana G Kirova, Jörg Mansfeld, Annette E Rünker, Gerd Kempermann

## Abstract

Intracellular redox states regulate the balance between stem cell maintenance and activation. Increased levels of reactive oxygen species (ROS) are linked to proliferation and lineage specification. In contrast to this general principle, we show that in the hippocampus of adult mice it is the quiescent neural stem cells (NSCs) that maintain the highest ROS levels (hiROS). Classifying NSCs based on intracellular ROS content identified subpopulations with distinct molecular profiles, corresponding to functional states. Shifts in ROS content primed cells for a subsequent transition of cellular state, with lower cellular ROS content marking activity and differentiation. Physical activity, a known physiological activator of adult hippocampal neurogenesis, recruited the quiescent hiROS NSCs into proliferation via a transient Nox2-dependent ROS surge. In the absence of Nox2, baseline neurogenesis was unaffected, but the activity-induced increase in proliferation disappeared. These results describe a novel mechanism linking the modulation of cellular ROS by behavioral cues to the maintenance and activation of adult NSCs.

**Highlights:**

- Quiescent adult hippocampal stem cells are characterized by high intracellular ROS
- Changes in intracellular ROS content precede changes in cellular state
- Acute physical activity recruits quiescent cells into active proliferation
- This recruitment is marked by a Nox2-dependent ROS spike in hiROS stem cells and represents an independent mode of cell cycle entry

**Graphical Abstract:** 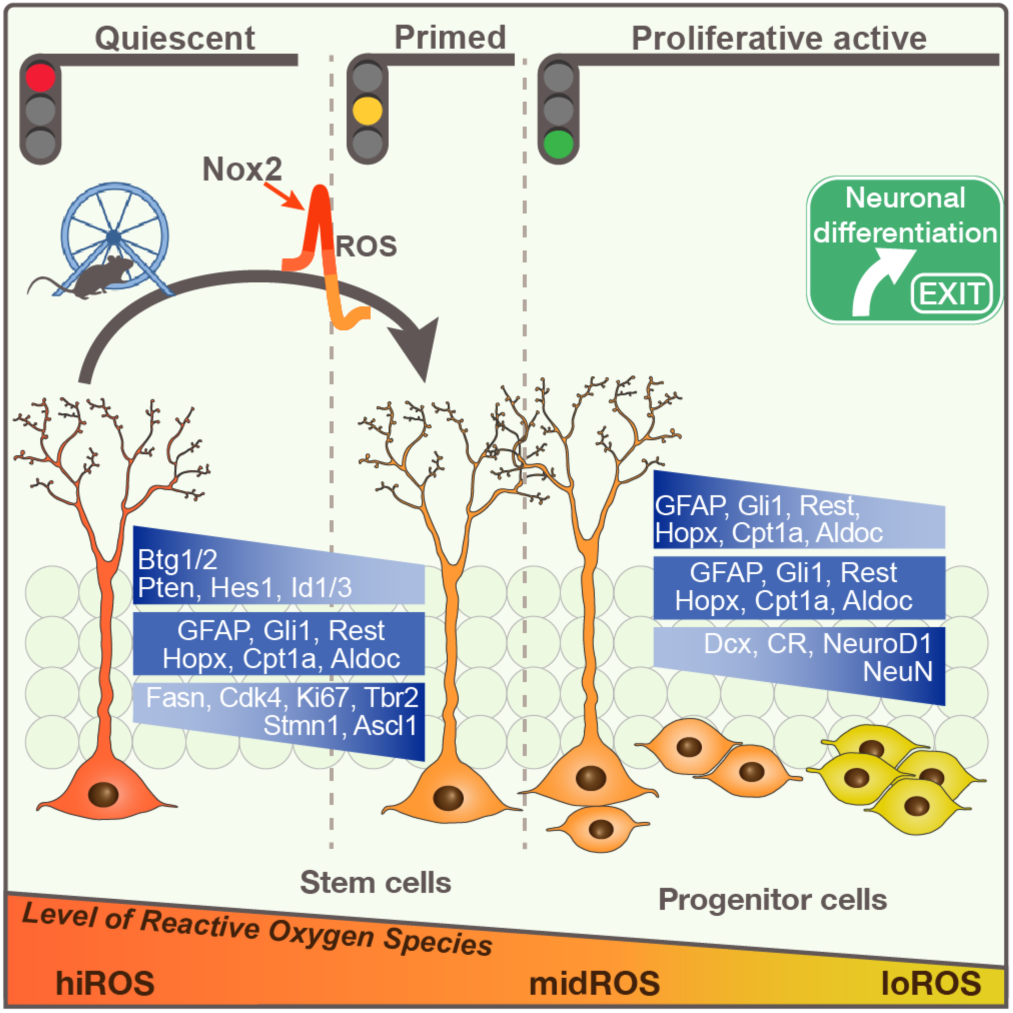

## Introduction

The dentate gyrus (DG) of the adult mammalian hippocampus is a unique structure that, unlike essentially all other parts of the brain, maintains adult neurogenesis of its principal neurons, the granule cells. The adult-generated neurons provide a particular type of plasticity, critical to the function of the DG (Garthe et al., 2009; Sahay et al., 2011). Histochemistry, in combination with state-of-the-art transcriptional profiling and longitudinal imaging, has provided foundational understanding of key aspects of adult neurogenesis (Bonaguidi et al., 2011; Encinas et al., 2011; Hochgerner et al., 2018; Pilz et al., 2018; Shin et al., 2015). Multipotent adult neural stem cells (NSCs / type-1 cells) with an elaborate radial glial morphology enter proliferation and, based on the symmetry of their division, undergo either self-renewal or generate transiently amplifying intermediate progenitor cells (type-2, type-3), which differentiate to yield post mitotic neurons that over weeks integrate into the local circuit (Kempermann et al., 2004). Different regulatory checkpoints within this process are highly responsive to physiological activity-driven stimulation. For example, physical activity and housing in enriched environments are strong pro-neurogenic cues in the DG (Kempermann et al., 1997; Kronenberg et al., 2003; van Praag et al., 1999). The responsiveness of adult neurogenesis to external stimuli poses particular challenges to the regulation of the NSC’s cell cycle entry, which needs to be tightly controlled yet sensitive to a wide range of distant cues. Although numerous mediating pathways have been described (Jang et al., 2013; Klempin et al., 2013; Lugert et al., 2010), it has remained elusive whether subsets of responsive cells exist within the known precursor cell populations, and whether differences in functional states might indicate how the actual activation takes place at the level of the NSCs themselves. We reasoned that an *a priori* differential enrichment of key molecular pathways might create functional subsets of NSCs and identified intracellular ROS content and cell-autonomous redox regulation as a key pathway delineating potentially unique functional states of the hippocampal precursor cells.

A cell’s redox potential, or cellular “oxidative stress”, is generally maintained by the balance between the generation and scavenging of reactive oxygen/nitrogen species (ROS/RNS). ROS is a product of incomplete reduction of molecular oxygen and intracellular ROS primarily exists as the highly reactive superoxide anion (O^-^_2_), which is enzymatically reduced to hydrogen peroxide (H_2_O_2_) and hydroxyl radicals (OH^-^) (Bigarella et al., 2014; Holmström and Finkel, 2014; Phaniendra et al., 2015). Sub-cellular compartments such as mitochondria, endoplasmic reticulum, lysosomes and the plasma membrane can generate ROS as a by-product of metabolic activity or through enzymatic activity of the Nox homologs (NADPH oxidizing enzymes), xanthine oxidase, cyclooxygenase etc. (Ye et al., 2015). The absolute intracellular ROS content, and its relative changes, can regulate the abundance of various redox sensors and thus can act as a physiological secondary messenger that integrates environmental cues and cell-autonomous signaling. Several lines of work have discussed this role for ROS fluctuations and oxidative stress in regulating stem cell fate in bone marrow, cardiac or induced pluripotent stem cells (iPSCs). In these cell types, lower ROS content generally marked self-renewal, with a more oxidized state marking the more differentiated states (Armstrong et al., 2010; Gurusamy et al., 2009; Smith et al., 2000).

A central role of redox status as a key functional determinant of “stemness” has been discussed in neural stem cell biology for many years, beginning with pioneering studies by Noble and colleagues (Noble et al., 2003; Noble et al., 2005). Despite intriguing findings for the second neurogenic niche in the adult mammalian brain, the subventricular zone (SVZ), supporting this general idea (Le Belle et al., 2011), this had never been addressed in the DG in any detail, even though DG and SVZ stem cells differ in very many aspects, including their susceptibility to become activated by behavioral stimuli. We here describe the differential enrichment of intracellular ROS, its unexpected specificity in regulating cellular state transition and mediating the activity-dependent recruitment of quiescent hippocampal NSCs into active proliferation.

## Results

### Physical activity recruits non-dividing, activatable stem cells into proliferation

Theoretically, an extrinsic stimulus such as physical activity (arguably the most elementary known pro-neurogenic behavioral stimulus) might act individually or synergistically, at different neurogenesis checkpoints (Overall et al., 2016). To characterize whether subsets of neural precursor cells (NPCs) have differential sensitivity to behavioral stimuli, we analyzed the responsiveness of proliferative and non-proliferative NPCs to acute de novo physical activity. We have previously shown that continuous running wheel access for 4 nights resulted in a significant increase in proliferating cells in the DG (Overall et al., 2013). We now investigated the dynamics preceding this activation by studying bouts of physical activity of variable duration (1, 2, 3, or 4 nights; Figure S1A). By comparing the number of dividing cells, marked by the incorporation of the thymidine analog BrdU during the S-phase of the cell cycle, we found that a minimum of 3 nights of physical activity were required to detect a statistically significant increase in the number of BrdU labeled cells (one-way ANOVA: *p* < 0.0001, *F*_(4,35)_ = 13.04; 0d: 1130 ± 125 cells vs. 3d: 1603 ± 93 cells, mean ± SEM, Dunnett: *p* = 0.01; increase by 42%; Figure 1A). At day 4, the proliferative pool had further increased to 183% of control values (4d: 2070 ± 142 cells; Dunnett: *p* < 0.0001; cf. Akers et al., 2014). Despite only being significant on the third day, there is a clear trend throughout the four-day period suggesting that the initial response may begin even earlier. This fact, coupled with differing statistical sensitivity due to varying group sizes etc., could possibly underlie the seemingly conflicting result presented in Steiner et al (2008).

**Figure 1.**
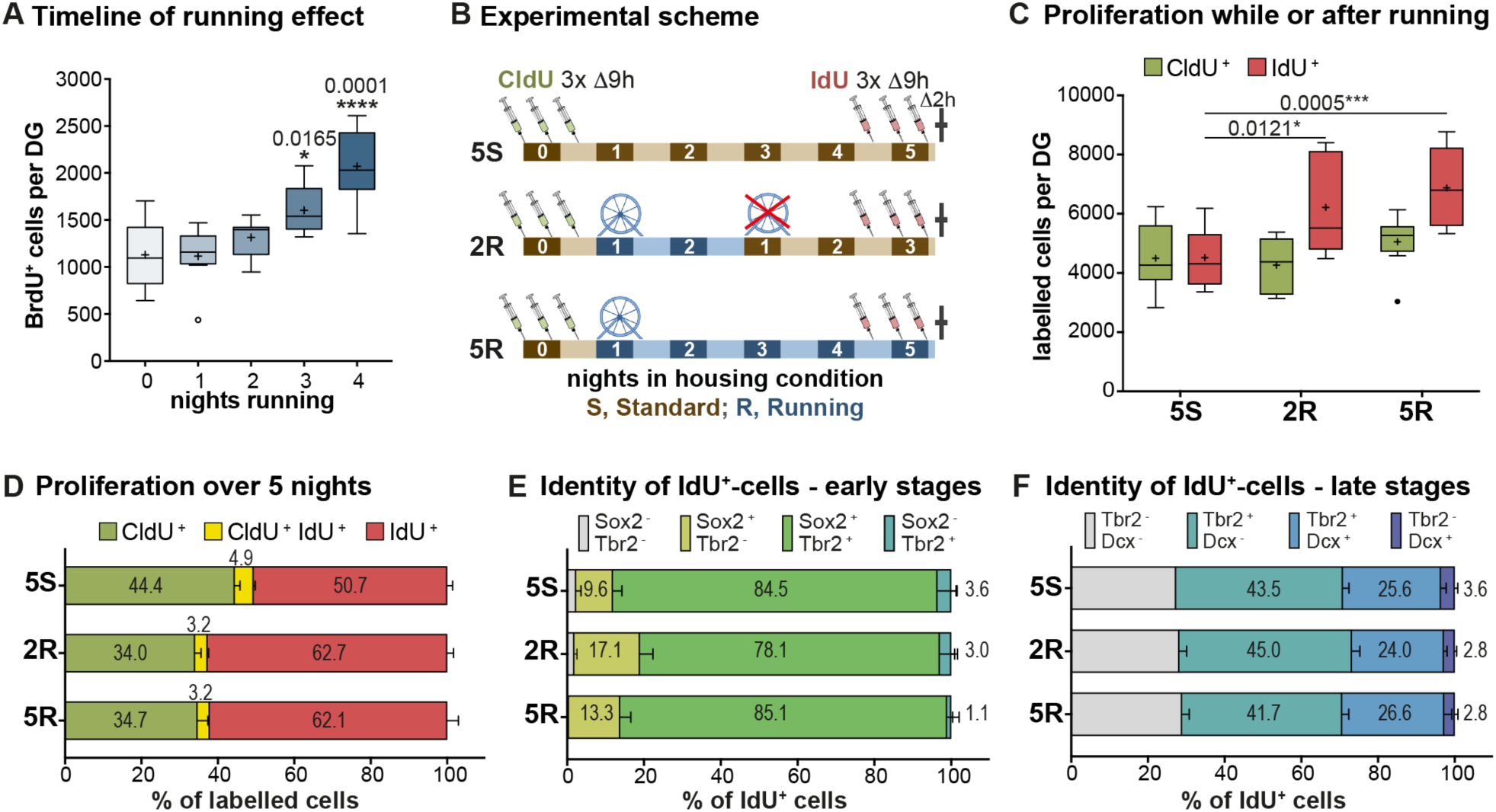
Physical activity stimulates quiescent cells into an invariable proliferation scheme. **(A)** At least 3 nights of physical activity are required to significantly increase the number of proliferating (BrdU^+^) cells in the DG. One-way ANOVA with Dunnett post-hoc test. **(B)** Experimental design for the dual thymidine labeling and running paradigm. **(C)** Although there is no change in baseline proliferating cells (CldU^+^), the number of IdU^+^ cells significantly increase in the 2R and 5R groups, indicating recruitment of quiescent cells. Two-way ANOVA with Dunnett post-hoc test. **(D)**The small proportion of CldU^+^IdU^+^ cells indicate that the majority of the proliferating cells had exited the cell cycle within the 4-day paradigm. **(E)** The proportion of the IdU^+^ cells that are “early” in the neurogenic progression (type-1 cells (Sox2^+^) or type-2 cells (Sox2^+^Tbr2^+^)). **(F)** The proportion of the IdU^+^ cells that are in “late” stages of neurogenic progression (type-2a cells (Tbr2^+^), type-2b cells (Tbr2^+^DCX^+^) or type-3 cells or post-mitotic immature neurons (DCX^+^)). All data represent mean ± SEM. * *p* < 0.05, *** *p* < 0.001, **** *p* < 0.0001.

To determine whether the increase in proliferation was primarily driven by an expansion of already dividing cells or the recruitment of quiescent cells into the proliferative cycle, we applied a dual thymidine analog (CldU, IdU) labelling paradigm (outlined in Figure 1B) (Shibui et al., 1989; modified from Brandt et al., 2012; Fischer et al., 2014). We first injected mice with CldU 3 times with an interval of 9 hours between injections (baseline). Animals were then individually housed, either with access to a running wheel (5R) or in standard housing conditions (5S), for a period of 5 nights (i.e. 4 days). To also detect a sustained pro-proliferative effect of an acute bout of physical activity, a third group of mice had running wheel access only for 2 nights (2R) before returning to standard conditions for the remaining 3 nights. Injections of IdU were used to mark the pool of dividing cells at the end of the stimulation period. Based on the number of IdU-positive cells, we found a strong pro-proliferative response, in the 5R group (6876 ± 474; increase by 52%; two-way ANOVA, *n* = 8: interaction: *p* = 0.04, *F*_(2,42) =_ 3.3, Dunnett: *p* = 0.0005, Figure 1C), compared to the 5S group (4517 ± 349). Interestingly, a statistically significant increase in the number of IdU-positive cells was also seen in the 2R group (6217 ± 581, increase by 38%, *p* = 0.01), suggesting that physical activity is not required as a continued stimulation but rather represents an activating event. In contrast, no significant differences were detected in the numbers of CldU-positive cells across the 3 groups (5S: 4496 ± 398; 2R: 4265 ± 319, *p* = 0.90; 5R: 5052 ± 328, *p* = 0.55). We noted that irrespective of the presence of the run stimulus, only a small proportion of CldU-positive cells was also positive for IdU (5S: 10.0 ± 0.9%; 2R: 8.7 ± 1.0%; 5R: 8.5 ± 0.4%; Figure 1D shows percentages of all labeled cells). Taken together, our results thus suggest that the increase in proliferation in response to exercise was due to an increased recruitment of previously non-proliferating cells.

To determine whether these quiescent cells, activated by physical activity, would predominantly expand the NSC stage, or instead progress to advanced stages of adult neurogenesis as do the cells proliferating in the absence of a run stimulus (Kronenberg et al., 2003), we phenotyped IdU-labeled cells using two different sets of antibodies: Sox2/Tbr2 to identify type-1 and type-2 cells (“early”, Figure 1E) and Tbr2/Dcx to identify type-2b and type-3 cells (“late”, Figure 1F). Under standard housing conditions, the majority of IdU-positive cells (84.5 ± 4.9%) were double positive for Sox2 and Tbr2 whereas 9.6 ± 2.4% were exclusively positive for Sox2 and 3.6 ± 1.6% showed only Dcx expression. Stimulation by physical activity (5R or 2R), did not significantly alter these proportions. These results suggest that physical activity stimulates a population of quiescent, yet activatable, precursor cells to enter proliferation without otherwise affecting the distribution among the neurogenic stages.

### Differential redox regulation of delineates sub-sets of precursor cells

To identify pathways involved in the activation of hippocampal neural stem cells, we compared the transcriptomic profiles of Nestin-GFP (Nes-GFP^+^) expressing precursor cells (Yamaguchi et al., 2000) from the DG to those from the SVZ. As SVZ cells do not show a neurogenic response to exercise (Figure S1B, C; Brown et al., 2003), we hypothesized that this comparison would reveal potential differentially enriched pathways responsible for maintaining a distinct population of activatable NSCs in the DG.

A principal component analysis (PCA) showed that the Nes-GFP^+^ cells from the two niches clustered distinctly (Figure 2A; see also correlation analyses in Figure S2). There were 4374 differentially expressed transcripts (30.2 % of all transcripts present) in the DG and 3826 (26.4 %) in the SVZ (Figure 2B). A Gene Ontology (GO) enrichment analysis of the differentially expressed genes showed that the top enriched pathways in the SVZ were related to ‘cell division,’ ‘transcriptional regulation,’ and ‘cell cycle’ (Figure 2C), which can be expected for proliferating cells. The enriched pathways of the DG, in contrast, did not show this association but were related to cellular responses to environmental modulations, and of these ‘redox regulation’ showed the greatest enrichment (p = 4^e-9^; Figure 2D).

**Figure 2.**
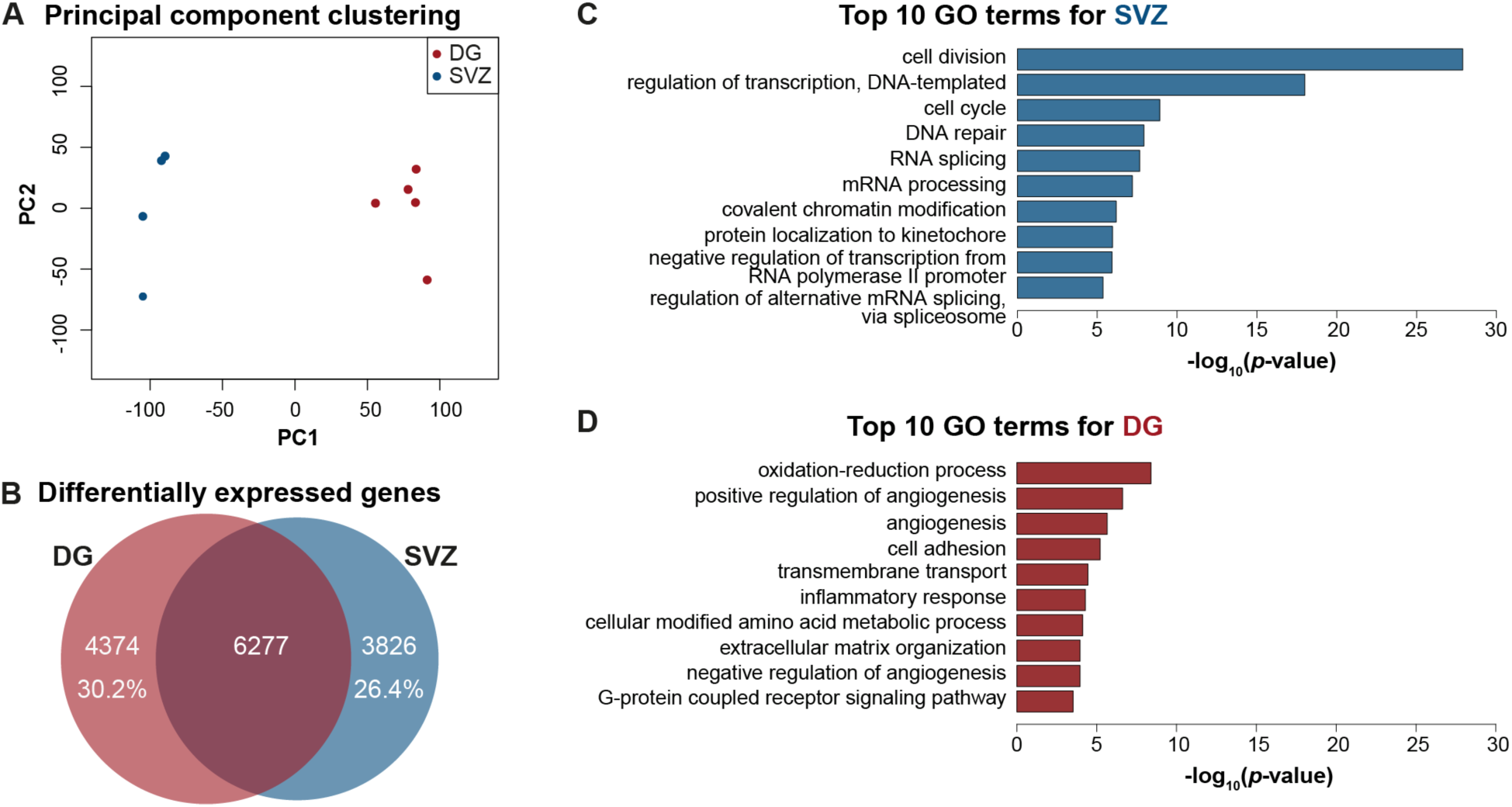
Nes-GFP^+^ cells of the DG and SVZ can be distinguished based in their differential regulation of ROS. **(A)** A principle component (PC) analysis showed that Nes-GFP^+^ samples from DG and SVZ cluster distinctly. **(B)** Approximately 30% and 26% of the detected transcripts are uniquely expressed in the precursor cells of the DG and SVZ respectively (signatures). **(C)** A Gene Ontology (GO) analysis of the signatures revealed that the most enriched pathways in the SVZ precursor cells were related to cell division and transcriptional regulation. **(D)** In contrast, redox regulation was identified as the most enriched pathway in the DG.

To validate whether ROS levels, and thereby the redox potential, are differentially maintained and regulated in the DG compared to the SVZ, we stained dissociated cells from both niches for total cellular ROS using the superoxide indicator dihydroethidium (DHE; Dikalov and Harrison, 2012; Zhou et al., 2016; Zielonka et al., 2008). We discovered that DG and SVZ cells isolated from the same animals had distinct ROS profiles (Figure 3A and E; gating strategy: Figure S3A-D). The SVZ cells, taken together, have 5.3-fold higher overall ROS levels than the DG cells (two-way ANOVA, *p* < 0.0001, *F*_(1,16)_ = 32.85; Tukey: *p* = 0.004; Figure 3I, left; compare also dotted lines in Figure 3C and G). We next applied the neurosphere bioassay to determine whether the cell-intrinsic ROS content of DG cells could predict their in vitro stemness (Reynolds and Weiss, 1992; Rietze and Reynolds, 2006). Indeed, almost all neurospheres (98.8%) were generated from the 17.8 ± 1.1% of FACS events with the highest ROS content (Figure S3F). This corresponds to an 8.2 ± 1.5 fold ROS-associated enrichment of neurosphere-forming cells relative to unsorted cells (one-sample t-test, vs. unstained, ROS^+^: adj. *p* = 0.01, ROS^-^: adj. *p* < 0.0002).

**Figure 3.**
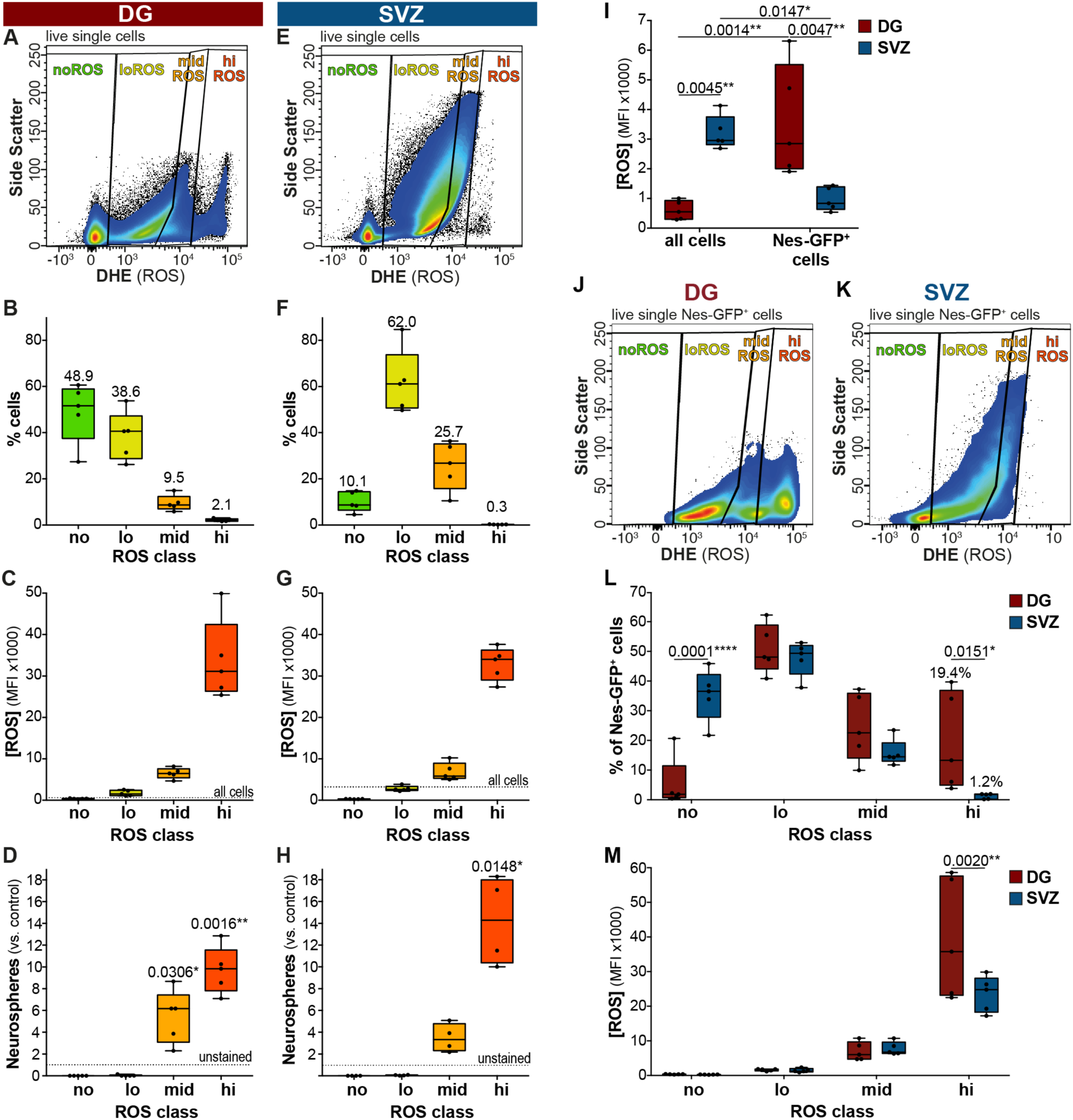
Nes-GFP^+^ cells can be classified into distinct functional subsets based on their cellular ROS content. Flow cytometric analysis revealed that the cells from the DG **(A)** and SVZ **(E)** have distinct ROS profiles. Plots showing the distribution **(B, F)** and total ROS content **(C, G)** of cells in each of the four manually gated ROS classes. Neurosphere formation is restricted to the midROS and hiROS class, with significantly more neurospheres formed in the hiROS class from both the DG **(D)** and SVZ **(H)**. One-sample t-tests with Bonferroni correction. **(I)** The total ROS content of all live cells in the SVZ is higher than that of the total DG cells. In contrast, the Nes-GFP^+^ cells of the DG contain significantly higher levels of cellular ROS that those of the SVZ. Two-way ANOVA with Tukey post-hoc test. FACS plots displaying the distribution of Nes-GFP^+^ cells from the DG **(J)** and SVZ **(K)** based on the total ROS content. **(L)** The DG and SVZ display unique distribution patterns across the four ROS classes, with the DG having significantly more Nes-GFP^+^ cells in the hiROS class. Two-way ANOVA with Sidak post-hoc test. **(M)** The Nes-GFP^+^ cells from the DG have significantly higher levels of cellular ROS than those isolated from the SVZ. Two-way ANOVA with Sidak post-hoc test. All data represent mean ± SEM. * *p* < 0.05, ** *p* < 0.01, **** *p* < 0.0001.

To further resolve the correlation between ROS content and the potential for neurosphere formation, we defined four distinct, non-overlapping classes of ROS levels (Figure 3A-C): hiROS (2.1 ± 0.3% events with highest ROS levels), midROS (next 9.5 ± 1.5% events), loROS (next 38.6 ± 4.7% events) and noROS (48.9 ± 5.8% events; Figure 3B). Applying the same gating strategy, in the SVZ (Figure 3F-G), we found that most cells were in the loROS (62.0 ± 6.2%) and midROS classes (25.7 ± 4.7%), with substantially fewer in the hiROS class (0.3 ± 0.0%). Interestingly from both the SVZ and DG, neurosphere-forming cells were restricted to the higher ROS classes (midROS and hiROS classes, which together constitute 26% and 11.6% of all sorted events from the SVZ and DG respectively), with the highest density of neurosphere-forming cells in the hiROS class compared to unstained, sorted cells (one-sample t-test, DG: adj. *p* = 0.002, SVZ: adj. *p* = 0.015; Figure 3D and H). Supplementing the growth medium with KCl or norepinephrine (NE), a treatment to boost neurosphere formation from DG precursors (Jhaveri et al., 2010; Walker et al., 2008), augmented neurosphere formation only from hiROS and midROS cells, with significantly more neurospheres induced in the hiROS class after KCl treatment (2.4 ± 0.7 fold; two-way ANOVA, *p* = 0.0029, *F*_(6,36)_ = 4.151; Dunnett: *p* < 0.0001; Figure S3G). Neither KCl nor NE elicited neurosphere formation from loROS and noROS classes. Based on these results and those by others (Le Belle et al., 2011), we conclude that cellular ROS content is an effective predictor of neurosphere-forming potential and that the stemness of adult neural precursor cells from both niches is related to their steady state cellular ROS content.

We hypothesized that a disproportionate concentration of Nes-GFP^+^ precursor cells in the higher ROS classes could explain the finding that these classes harbor the neurosphere-forming cells. We found that the ROS content of Nes-GFP^+^ cells of the DG was significantly greater than that of all isolated cells (3.7-fold; Tukey, *p* = 0.001; Figure 3I). Surprisingly, however, the distribution analysis of Nes-GFP^+^ cells of the DG into the four ROS classes (Figure 3J; gating: Figure S3A-E) revealed the following characteristic pattern: 50% of the Nes-GFP^+^ cells were negative for or low in ROS (noROS: 5.3 ± 3.9% of all Nes-GFP^+^ cells; loROS: 50.8 ± 3.7%), thus representing sub-populations that are not competent to form neurospheres. An additional 24.5 ± 5.1% of Nes-GFP^+^ cells localized to the midROS and 19.4 ± 7.4% to the hiROS class (Figure 3L). In contrast to the DG, the Nes-GFP^+^ cells of the SVZ had significantly lower absolute ROS levels than the entirety of cells isolated (Tukey, *p* = 0.0047; Figure 3I). In terms of their relative distribution among ROS classes, in the SVZ (Figure 3K) significantly more precursor cells segregated to the noROS class (35.3 ± 3.9%; two-way ANOVA, *p* < 0.0001, *F*_(3,32)_ = 13.0; Sidak, *p* < 0.0001), and significant fewer (1.22 ± 0.4%, p = 0.01) were found in hiROS compared to the DG (Figure 3L). In summary this suggests that, first, precursor cells of the DG are maintained at, and marked by, a higher cellular ROS content compared to the surrounding other DG cells; and, second, that ROS content resolves the precursor pools of DG and SVZ into distinct functional classes and, third, compared to the SVZ, the Nes-GFP^+^ hiROS population is substantially enriched in the DG.

### Nes-GFP^+^ cells of the different ROS classes have distinct molecular profiles

To verify if intracellular ROS-based clustering identifies cell populations with truly distinct functional states, we isolated and profiled the total RNA from Nes-GFP^+^ cells of the DG sorted into loROS, midROS and hiROS classes by means of next generation sequencing (NGS). As only a very small proportion of Nes-GFP^+^ cells clustered into the noROS fraction (5.26 ± 3.87%), we omitted this group from the following analysis. The Nes-GFP^+^ cells in the different classes of cellular ROS content also had distinct transcription profiles, as shown in a principal component analysis (PCA; PC1: 41% and PC2: 13%; Figure 4A). As evident from the PCA plot and correlative analysis of the different biological samples (Figure S4A-C), we found tight clustering of the expression profiles within a class: 47.7% of the expressed transcripts segregated based on ROS content with 36.1% of transcripts being specific to cells of only one of the ROS classes. We could identify 1828 transcripts (12.1% of all expressed transcripts) specific for the hiROS class, 1043 transcripts (6.9%) for the midROS class and 2589 transcripts (17.1%) for the loROS class (Figure 4B). Additionally, 684 transcripts (4.5%) were co-enriched between high and midROS class and 1050 transcripts (6.9%) among the mid and loROS classes. Interestingly, only 25 transcripts (0.2%) were co-enriched in the Nes-GFP^+^ cells of high and loROS, indicating that these ROS classes are the most distinct (Figure 4B). Transcripts from Nes-GFP^+^ cells that were specific to the hiROS class were more likely to be expressed in the DG than the SVZ, whereas transcripts exclusive to the Nes-GFP^+^ cells with loROS were more likely to be found in the SVZ (Figure 4C). To better resolve functional heterogeneity, we looked at expression of known transcripts that mark cellular processes such as neurogenesis, cell cycle, and, as a control, ROS regulation. As expected, across the groups from hiROS to loROS we found a decline in the expression of transcripts considered to be essential for maintaining higher intracellular levels of ROS. We also found a concomitant increase in markers of active cell cycle and neurogenic commitment (Figure 4D). Nes-GFP^+^ cells of high and midROS expressed general neural stem cell markers such as Nestin, Gfap, Notch2, Lpar1 (Walker et al., 2016; complete list of queried genes in Supplementary File 6). There was negligible expression of Eomes (Tbr2; Hodge et al., 2012), E2f1, a potent cell cycle regulator (Black et al., 2005; Cooper-Kuhn et al., 2002), and no enrichment for proliferation markers (Pcna, Mki67, MCMs and others). The level of expression for cell cycle inhibitors Cdkn1a (Pechnick et al., 2008) and Pten (Bonaguidi et al., 2011; Hill and Wu, 2009), which are critical in controlling proliferation and cell cycle exit of stem cells, was highest in the hiROS class. The hiROS cells also had comparatively high levels of the Notch ligand Jag1, as well as Notch1 and 4, and Sirt1, which might be seen as distinct cellular sensors of the microenvironment. Genes that reflect the ability of the precursor cell to enter proliferation such as Gli1, Id4 (Bedford et al., 2005; Boareto et al., 2017), Yap1, Runx1, and Foxo1 were highest in cells of the hiROS and midROS classes.

**Figure 4.**
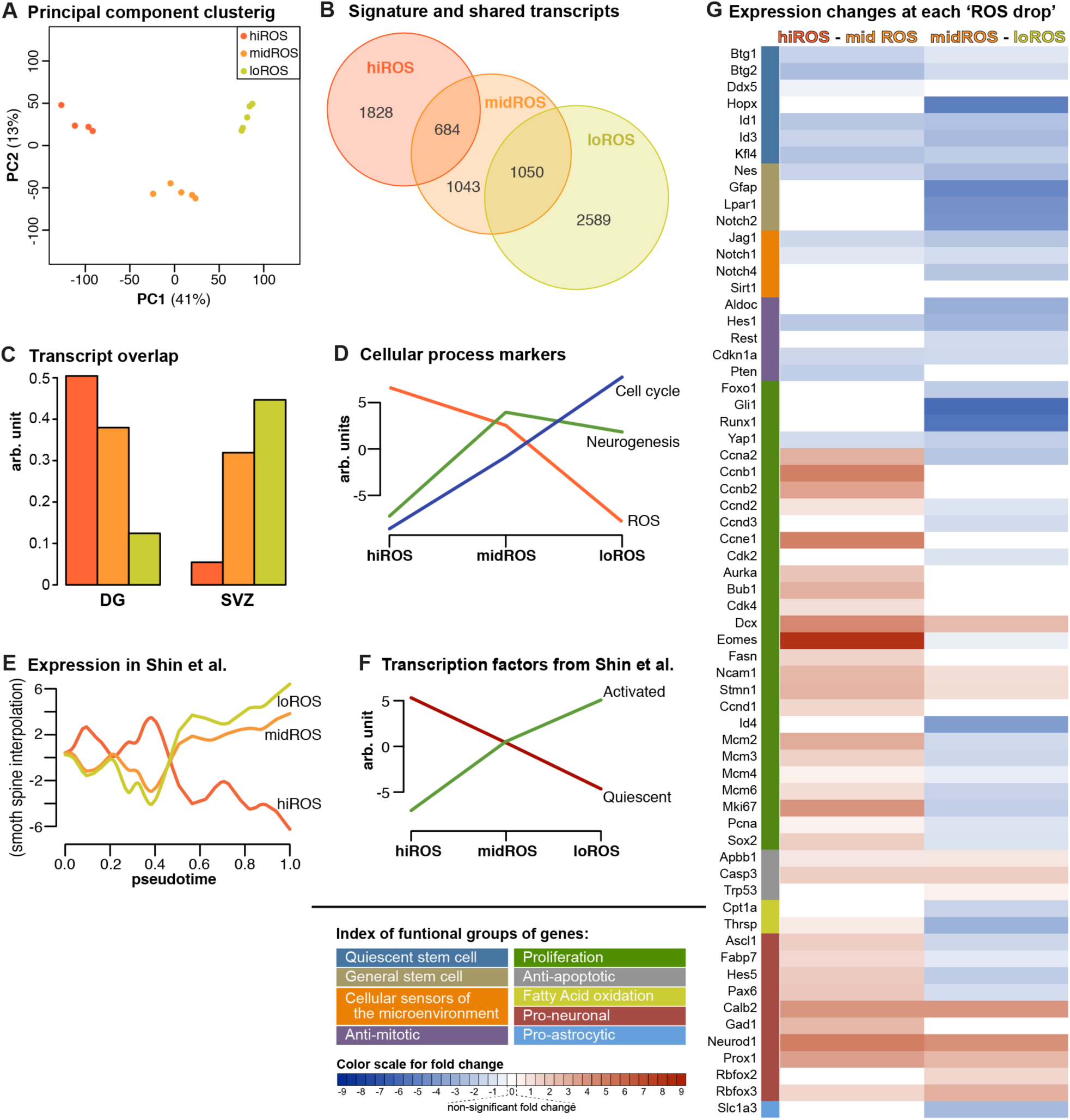
ROS levels decrease along the neurogenic trajectory. **(A)** Principal component (PC) analysis shows that samples from the different ROS groups have distinct transcriptional profiles. **(B)** Venn diagram showing the extent of differential gene expression between the Nes-GFP^+^ cells of the different ROS classes. Note that 0.2% transcripts are co-enriched in hiROS and loROS classes, and are omitted in this panel. **(C)** Transcripts uniquely enriched in hiROS cells (signature) are more likely to be expressed in the DG than the SVZ, while loROS signature transcripts are more likely to be found in the SVZ. **(D)** Decreasing ROS is associated with an increase in cell cycle activity and neural differentiation, and an expected decrease in ROS marker transcript expression. **(E)** In a pseudotime trajectory as defined by Shin et al. (2015), Nestin^+^ cells are arranged along the *x* axis from quiescent stem cells to more mature neuronally-committed cells. Plotting expression of the ROS groups in these cells showed higher expression of hiROS transcripts in cells early in pseudotime, with later peaks in expression in the lower ROS groups. **(F)** Decreasing ROS content is associated with increased expression of transcription factors associated with activated stem cells (as identified by Shin et al. 2015) as well as a decrease in expression of quiescent stem cell transcription factors. Panels D-F were plotted using the first principal component of the transcripts associated with each gene list (and are thus on an arbitrary *y* scale centred on 0); see Supplementary Methods for details. **(G)** Heatmap showing the changes in expression of specific genes at the high to midROS transition (left) and mid to loROS transition (right). Colour scheme indicates significance of change (white indicates no significant change) and the degree of fold change.

The Nes-GFP^+^ cells of the midROS class showed a complex transcript profile. On one hand, similar to the hiROS cells, cells in this class were characterized by expression of genes such as Rest (Gao et al., 2011), Hes1 (Hatakeyama et al., 2004) and Aldoc (Shin et al., 2015), which negatively control cell cycle entry and maintenance. Furthermore, Thrsp (Spot14; Knobloch et al., 2014) and Cpt1a (Knobloch et al., 2017), genes essential for fatty acid oxidation, a critical pathway associated with precursor cell quiescence and a switch to active proliferation, were uniquely enriched in the midROS class. On the other hand, this class was also characterized by the highest levels of expression of proliferation markers such as Pcna, Mki67 and Mcm2/3/4/6. We detected cyclins/CDKs required for G1/S transition such as E2f1, Ccnd2/3, Ccne1, Cdk2, Ccna2 and Ccnb1/2 as well as cell cycle promoters Bub1 and Aurka. The midROS cells had the highest levels of Sox1, 2 and 9 and expressed both pro-neural genes such as Ascl1 (Andersen et al., 2014; Kim et al., 2011), Hes5 (Lugert et al., 2010), Pax6, Fabp7 (BLBP; Giachino et al., 2014) and genes of the astrocytic lineage such as Slc1a3 (GLAST; Codega et al., 2014; Doetsch, 2003) highlighting that these cells of the DG, with moderate levels of ROS levels, are actively engaged in, or poised for, the growth phase of the cell cycle. Genes like Dcx, Stmn1, Fasn (Knobloch et al., 2012), Eomes (Tbr2), Cdk4 and Ncam1, which are indicative of a high degree of commitment to proliferation were co-enriched in Nes-GFP^+^ cells of both the midROS and loROS classes. The Nes-GFP^+^ cells of the loROS class were uniquely enriched for transcripts marking neuronal differentiation including advanced stages, such as Calb2 (calretinin), Rbfox3 (NeuN), Rbfox2, Prox1, Neurod1 and Gad1. Additionally, SoxC (Sox4 and Sox11) were enriched together only in the loROS class, along with Nfib and Tcf4. The cells of the loROS class were also enriched for anti-apoptotic and DNA damage checkpoint markers such as Trp53 (p53), Casp3 and Apbb1. Taken together, these results suggest that at a reduced intracellular ROS content, the Nes-GFP^+^ cells are committed to proliferation and neuronal lineage (see expression heat map of 175 neurogenesis-related transcripts, Figure S4G).

To corroborate our assertion that the segregation of Nes-GFP^+^ cells into sub-classes based on decreasing cellular ROS content indeed represents a trajectory from quiescence to neuronal commitment, we compared our findings to an existing sub-clustering of DG precursor cells. Using Nes-CFP^nuc^ mice, Shin and co-workers had provided comprehensive single cell transcriptomic data of the DG precursor cell population and had aligned the cells along a reconstructed developmental trajectory (‘pseudotime’) ranging from quiescent to actively dividing cells (Shin et al., 2015). We investigated the expression of the genes uniquely characterizing each of our ROS groups (‘signature’ genes) in the dataset from Shin et al. (see Methods for details). The expression of genes associated with both the hiROS and midROS classes were more strongly expressed early in pseudotime, while the genes associated with the loROS class were expressed more strongly later in the reconstructed trajectory (Figure 4E). We next analyzed the expression of 82 transcription factors (TFs), attributed by Shin et al. to the quiescent and active proliferative status of precursor cells, in our three ROS classes. Interestingly, we found a clear similarity between TFs regulating quiescence and genes characterizing the hiROS cells, a relationship which inverts with the drop in ROS content. Conversely, TFs associated with active proliferation were almost absent in hiROS cells but expression increased in cells with lower ROS content (Figure 4F).

### Changes in intracellular ROS content precede cell fate changes

From our re-analysis of the data from Shin et al., it appeared that the transcripts necessary for ROS production would be gradually downregulated along the pseudotime trajectory of Nes-GFP^+^ cells. However, a substantial drop in cellular ROS content is observed between Nes-GFP^+^ cells of the hiROS and midROS classes (~5-fold reduction in ROS content; see also Figure 3A, C). To investigate expression changes at this transition more closely, we performed a pairwise comparison of the expression profiles of the hiROS and midROS classes using a different statistical model (see Methods). We found that the high to mid ROS transition was marked by with a downregulation of cell cycle inhibitors (Pten, Btg1/2, Hes1, Cdkn1a, Klf4, Ddx5), and an upregulation of markers for active proliferation (Sox2, Eomes, E2f1, Pcna, Fasn, Dcx, Stmn1, Hes5, Fabp7, Prox1, Ccnd1/2, Cdk4, Sirt2). However, we detected unchanged expression of many radial glial markers (Gfap, Egfr, Gli1) and quiescence markers such as Aldoc, Hopx, Thrsp, Cpt1a, Rest (among others; Figure 4G, see Supplementary File 5 for complete gene list and GO terms associated with the ROS shift). These results suggest that although the ROS transition is associated with expression of key genes necessary to enter the active growth cycle, this switch in ROS content precedes downregulation of all genes maintaining the quiescent state, indicating that ROS content would delineate a functional rather than molecular state.

A second drop in ROS levels (~2 fold) occurred between the two putative proliferative fractions, midROS and loROS, and was marked by a decrease in classical NSC markers (Gfap, Egfr, Nes, Slc1a3, Prom1, Notch2, Lpar1), markers for active proliferation (Sox2, Eomes, Pcna, Mki67, Mcm2/3/4/6, Hes5, Pax6), and astrocytic markers (Fabp7). Concurrently, quiescent markers (Hopx, Cpt1a, Thrsp, Aldoc) were downregulated, while Stmn1 was upregulated together with redox sensors such as Foxo3 and Trp53 (Figure 4G). At the same time, this shift in ROS content from moderate to lower levels marked an increase in markers of neuronal commitment (Rbfox2/3, Neurod1, Ncam1, Prox1, Calb2) (see Supplementary File 5 for complete gene list and GO terms associated with ROS shift), which, as we postulate, marks the cellular progression from transient amplification to a neuroblast-like identity. Taken together these findings suggest that ROS content delineates functional states among NSCs. This finding goes beyond previous characterizations and refines our model of the hippocampal NSC niche. We propose that stem cell ‘activatability’ might be the functional equivalent of the hiROS state.

### Nes-GFP^+^ cells of the hiROS class specifically respond to physical activity

As a next step, we sought to link these data to our initial observation that the pro-neurogenic response elicited by acute physical activity involves the recruitment of quiescent cells into active proliferation. We reasoned that if hiROS Nes-GFP^+^ cells indeed represented the activatable quiescent NSC pool, then changes in ROS levels in these cells should precede the overall increase in proliferative cells in response to physical activity. We subjected Nes-GFP-reporter mice to two such physical activity paradigms, first, an acute, suboptimal bout of one night of wheel access, which is not sufficient to elicit a measurable increase in proliferation (Figure 1A), and the exposure to four nights of wheel access, which elicits the proliferative response. Analyzing the NSC ROS profiles under these conditions, we found that in response to one night of running, the distribution of Nes-GFP^+^ cells among ROS classes was not altered in comparison to control conditions (Figure 5A). However, there was a robust and significant surge in the intracellular ROS content exclusively in the Nes-GFP^+^ cells of the hiROS class (increase to 152.6 ± 25.3%, two-way ANOVA, *p* = 0.02, *F* (_6,64_) = 2.703; Tukey: hiROS Std vs. 1dRun *p* = 0.001; Figure 5B). This acute ROS response had the characteristics of a spike, since the ROS content returned to control levels after four nights of physical activity (Tukey: Std vs. 4dRun *p* = 0.44 ns; 1dRun vs. 4dRun *p* < 0.0001; Figure 5B). However, at this later timepoint, which follows the acutely induced fluctuations in ROS content, the distribution of Nes-GFP^+^ cells into ROS classes had distinctly changed compared to standard housed controls. We detected an increase by a third (from 50.8 to 67.2% of all Nes-GFP^+^ cells) in the proportion of Nes-GFP^+^ cells in the proliferative and neuronally committed loROS class (Tukey, Std vs. 4dRun for loROS, *p* = 0.03; Figure 5A). In the SVZ, the Nes-GFP^+^ cells of the hiROS class showed a similar ROS spike, however this class of precursor cells represents a fraction of less than 1% of the Nes-GFP^+^ cells, which is presumably far too small to translate into a meaningful neurogenic response (Figure S5A, B). Taken together, these results indicate that Nes-GFP^+^ cells of the DG, with respect to their redox state, show distinct temporal responses to physical activity. An increased ROS level of hiROS Nes-GFP^+^ cells, which constitute about 15-20% of Nes-GFP^+^ cells in the DG, is the “first response” to acute physical activity, which is followed by their redistribution among ROS classes over the course of 4 days.

**Figure 5.**
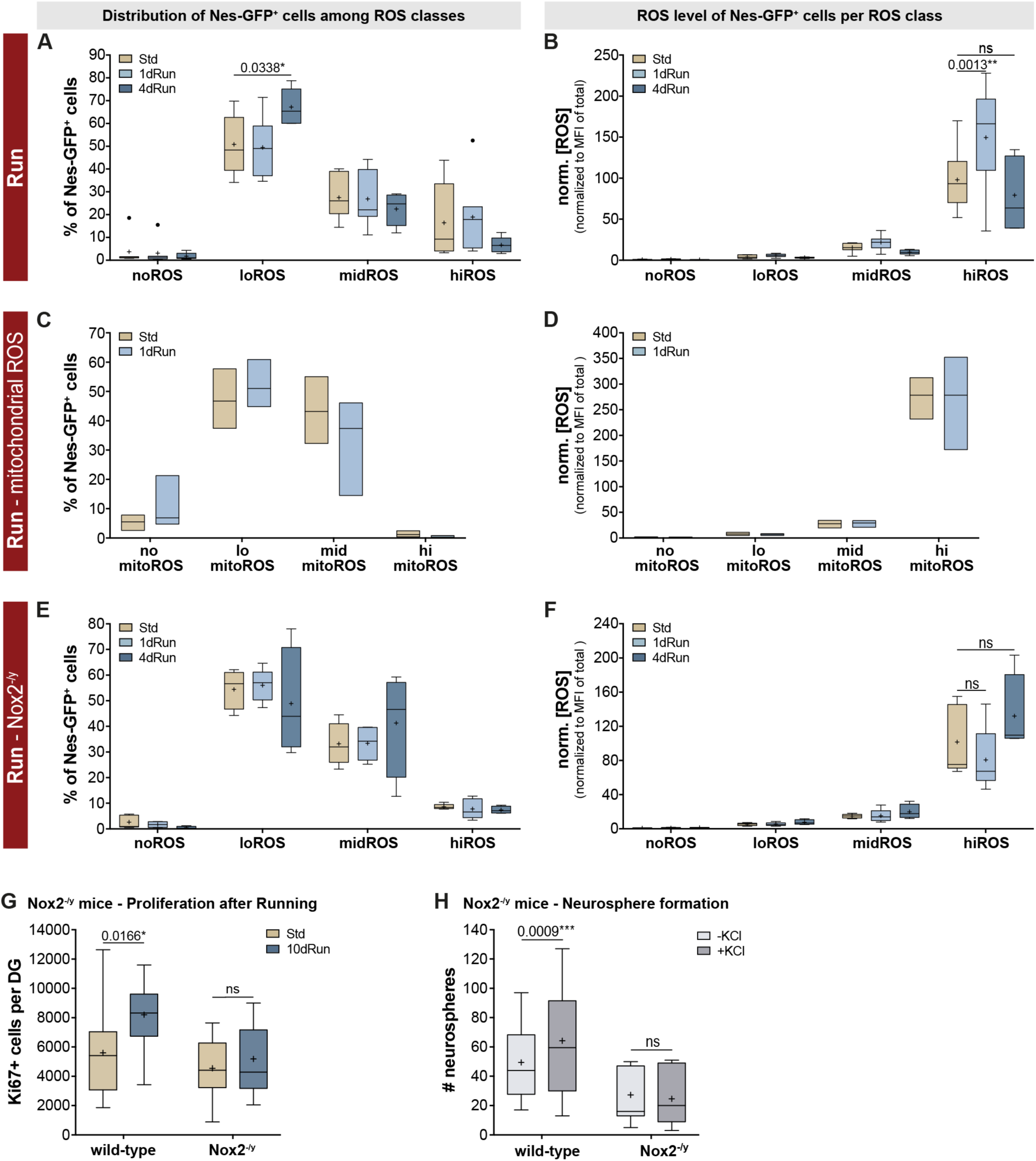
Nox2-mediated ROS fluctuations in the hi-ROS fraction of Nes-GFP+ cells are critical for physical activity-mediated precursor activation. **(A)** Although 1 day of physical activity did not alter the distribution of Nes-GFP^+^ cells from the DG across the four ROS classes, a significantly increased proportion of Nes-GFP^+^ cells could be found within the loROS class following 4 days of physical activity. **(B)** 1 night of wheel running however, caused a significant surge in ROS in the Nes-GFP^+^ cells of the hiROS cluster, which returned to levels similar to standard housed animals after 4 days of physical activity. Two-way ANOVA with Tukey post-hoc test. **(C)** Changes in ROS content in response to exercise are not driven by mitochondrial activity, with no change observed in either the distribution of Nes-GFP^+^ cells into the four mitoROS classes or mitochondrial ROS content **(D). (E)** One night of physical activity did not change the distribution pattern of Nes-GFP^+^ cells in the different ROS classes in Nox2 mutants, additionally, four nights of physical activity did not result in a redistribution of Nes-GFP^+^ cells into the ROS classes. **(F)** In contrast to the ROS surge observed in the wildtype Nes-GFP^+^ cells of the hiROS class, no changes in ROS levels in corresponding class of the Nox2 mutants was observed. Two-way ANOVA with Tukey post-hoc test. **(G)** Although the Nox2 mutants had similar baseline proliferation rates, 10 nights of physical activity was able to increase the number of Ki67^+^ cells in the WT but not the Nox2 mutant animals. Two-way ANOVA with Sidak post-hoc test. **(H)** Furthermore, while the number of neurospheres that were generated from the DG of wildtype and Nox2 mutant mice was not significantly different, the addition of depolarizing levels of KCl to the medium was only able to increase the neurosphere number in the wildtype animals. Two-way ANOVA with Sidak post-hoc test. All data represent mean ± SEM. * *p* < 0.05, ** *p* < 0.01, *** *p* < 0.001.

### Changes in ROS content are not driven by mitochondrial activity

Mitochondrial respiration is the major source of intracellular ROS (Jensen, 1966; Zorov et al., 2014) and mitochondrial maturation is critically linked to proliferation and progression through the developmental stages of neurogenesis (Beckervordersandforth et al., 2017). We therefore asked whether the acute ROS surge seen after one night of running was driven by mitochondria-generated ROS (mitoROS) production. To address this question, we used Mitosox red (Robinson et al., 2006), a mitochondria-targeting version of the DHE dye that was used for most of our aforementioned experiments, in order to specifically analyze the distribution pattern of Nes-GFP^+^ cells according to their mitoROS levels and to observe potential mitoROS fluctuations after one night of running. Classifying DG cells into four mitoROS classes by applying a similar gating strategy to that described for cellular ROS, we found fewer Nes-GFP^+^ cells in the hiROS class (1.15 ± 0.64%) than after staining for cellular ROS (16.40 ± 6.04%; two-way ANOVA, *p* = 0.049, *F*_(3,32)_ = 2.91; cellular vs. mitoROS not different). Further, comparing standard housed mice with animals running for one night, we did not find any changes in distribution of Nes-GFP^+^-cells and, more importantly, there was no discernable increase in mitoROS (Figure 5C and D), suggesting alternate sources for the ROS surge after a run stimulus.

The other source of intracellular ROS besides mitochondria is the NADPH oxidase enzyme complex (Nox; Le Belle et al., 2011; Dickinson et al., 2011). We therefore analyzed Nes-GFP^+^ cells of the different ROS classes for their transcript expression of Nox isoforms. The transcripts for plasma membrane-bound Nox2 (Cybb) and p22phox (Cyba) and for the cytosolic P67phox (Ncf2) were the highest in Nes-GFP^+^ cells of the hiROS class, while those of other subunits (Ncf4, Rac1/2) were maintained in Nes-GFP^+^ cells of the hiROS and midROS classes. Interestingly the transcript levels of Cyba, Cybb and Ncf2 decreased significantly at the ROS transition from hiROS to midROS, with even lower levels in cells of the loROS class (Figure S4E). This strongly suggested to us that the Nox2 complex might, on demand, produce ROS in the Nes-GFP^+^ cells of the hiROS class. We thus crossed Nes-GFP-reporter into Nox2-deficient mice (see methods for complete genotype; Nox2^-/y^ males; (Pollock et al., 1995) and analyzed the ROS profiles in this mouse model. Under baseline conditions, we detected similar distributions of Nes-GFP^+^ cells among ROS classes (Figure S5C, two-way ANOVA; genotype: *p* = 0.97, *F*_(1,40)_ < 0.01) with similar ROS profiles (Figure S5D; two-way ANOVA; genotype: *p* = 0.85, *F*_(1,40)_ = 0.04) from reporter Nes-GFP and Nes-GFP::Nox2^-/y^ mice. Analogous to the findings in Nes-GFP^+^ animals (Figure 5A, B), in the Nox2 mutants, one night of physical activity did not change the distribution of Nes-GFP^+^ cells in the different ROS classes (Figure 5E). Nevertheless, instead of the ROS surge seen in the wildtype Nes-GFP^+^ cells of the hiROS class, the ROS content in the Nox2 mutants decreased to 79.3 ± 16.8% in response to the stimulus (two-way ANOVA; housing: *p* = 0.13, *F*_(2,44)_ = 2.17; Figure 5F). Furthermore, four nights of physical activity did not result in a redistribution of Nes-GFP^+^ cells among ROS classes or changes in intracellular ROS content (Figure 5E and 5F). These results suggest that the ROS surge and the subsequent change in distribution of Nes-GFP^+^ cells into the ROS clusters, elicited by acute physical activity, is absent in Nox2 deficient mice and, thus, must be driven by Nox2 activity.

We confirmed that the Nox2-mediated ROS surge is indeed essential for a pro-neurogenic response by quantifying Ki67^+^ cells in the DG of wildtype and Nox2 mutants, under standard conditions and after 10 nights of physical activity. Under baseline conditions, Nox2-deficient animals had similar proliferation rates, as measured by Ki67^+^ cell numbers, compared to wildtype littermates (two-way ANOVA; genotype: *p* = 0.004, *F*_(1,46)_ = 9.235, housing: *p* = 0.012, *F*_(1,46)_ = 5.879; Tukey post-hoc: wt-Std vs. Nox2^-/y^-Std *p* = 0.70; Figure 5G). However, while physical activity robustly increased Ki67^+^ cells in wild-type animals (*p* = 0.02), Nox2^-/y^ mice exhibited no such increase (*p* = 0.91; Figure 5G). Additionally, the ex vivo neurosphere assay revealed that addition of KCl to the culture medium did not have any stimulatory effect on neurosphere formation from Nox2 mutant animals (two-way ANOVA, matched within genotypes; interaction: *p* = 0.0019, *F*_(1,19)_ = 12.97; Sidak: wt *p* = 0.0009, Nox2^-/y^ *p* = 0.68; Figure 5H). Taken together, these results show that Nox2-dependency is a distinguishing feature between baseline control of adult neurogenesis and the acute activation through the pro-proliferative stimulus of physical activity.

## Discussion

We here describe that functionally distinct sub-populations of adult neural stem cells can be defined by the intracellular ROS content and its activity-dependent fluctuations. With the help of transcriptional profiling we show that a classification based on ROS content aligns with other categorizations based on gene expression and identifies neural precursors with distinct cellular states. We show that conditions in the adult hippocampus deviate from what has been described for other systems (Armstrong et al., 2010; Ludin et al., 2014) in that in the DG intracellular ROS content decreases with increased commitment to proliferation. Cell-intrinsic shifts in ROS in bona-fide quiescent cells that are already high in intracellular ROS precede cell cycle entry and neurogenesis. Further, a specific Nox2-dependent ROS pulse in this population of cells is required for the activity-dependent stimulation of adult neurogenesis. These data suggest that ROS signaling and redox state are indeed defining features of activatable NSCs in the adult brain. These functionally delineated cells are highly enriched in the DG, which in contrast to the SVZ exhibits activatability by physical activity (Brown et al., 2003).

The point of origin of our investigation was the observation, here shown in Figure 1, that an acute bout of physical activity, a well-described physiological stimulus of precursor cell proliferation and neurogenesis in the adult hippocampus (Farioli-Vecchioli et al., 2014; Praag et al., 1999, Overall et al., 2013) functions as an activation event to recruit non-dividing cells into the cell cycle. Cells that were engaged in active cell cycle do not respond to acute physical activity. A difference between baseline control and activity-dependent regulation has been suggested before (Kempermann, 2011). Several previous studies have identified factors such as Igf1 (Trejo et al., 2001), VEGF (Fabel et al., 2003), endocannabinoids (Hill et al., 2010), serotonin (Klempin et al., 2013) and T cells (Walker et al., 2018), as specific mediators for distinct stages of adult hippocampal neurogenesis under standard housing and physical activity conditions. The key novelty here is that now such selective effects can be broken down temporally and to the level of a distinct subset of responsive cells, characterized by ROS content and the expression of related transcripts.

We used Nestin-GFP as a reporter for precursor cells. Although the proportional distribution of stem and progenitor cells positive for Nestin differs between the two niches, DG and SVZ, this is the broadest single marker recognizing early cell stages of neurogenesis. We show that a high level of intracellular ROS characterizes those NSCs from both adult neurogenic niches that harbor the highest ex vivo stemness. Based on our results, we postulate that hippocampal NSCs, routinely identified by the expression of GFAP, Nestin, Gli1, EGFR, and Hopx (among other markers), are functionally heterogeneous and contain a reproducible subset of cells that are identified by high cellular ROS content and which can be characterized by (1) the restricted gene expression of bona fide quiescent stem cell markers (Btg1/2, Klf4, Id1/3 and others), (2) the complete absence of the transcript for Eomes (Tbr2) and (3) three-fold lower levels of Mki67 and E2f1 transcripts as markers for proliferation.

We propose that through a drastic shift in cellular ROS content these quiescent cells convert to a poised-yet-not-yet-proliferative subset of NSCs that have only moderate cellular ROS content (midROS Nes-GFP^+^ pool) and which continue to maintain the expression of genes implicated in cell cycle repression or associated with quiescence, such as Hopx, Cpt1a, Thrsp (Spot14), Aldoc and Rest at the same levels as in the hiROS population. Based on our results we argue, therefore, that changes in ROS may precede other metabolic changes, such as the shift from fatty acid oxidation to lipogenesis (Knobloch et al., 2012) or the shift to oxidative phosphorylation as the major energy source in the NSCs (Beckervordersandforth et al., 2017; Zheng et al., 2016), both of which have been shown to be necessary for entry into the active growth cycle. The poised status of these Nes-GFP^+^ midROS cells, however, can be concluded from a marked reduction in transcripts for key cell cycle inhibitors such as Pten and Cdkn1a, and an upregulation of transcripts necessary for proliferation such as Ascl1, Eomes, Mki67, E2f1, Pcna and Mcm2/3/4/6. Cells seemingly preserve these moderate cellular ROS levels as they traverse through transient amplification (corresponding to the type-2a stage), a phase marked by upregulation of Fasn, Stmn1, Ccnd1/2 and Cdk4. We further speculate that the second reproducible ROS shift leading to Nestin-GFP^+^ cells of the loROS class marks the transition into a “neuroblast”-like identity (type-2b/type-3), marked by reduced proliferation and a more neuronal profile. Our hypothesis that ROS changes precede neurogenic progression aligns well with the trajectories proposed by Shin et al. based on their pseudotime analysis of single cell RNAseq data (Shin et al., 2015).

Although from our results we cannot yet predict the conversion rates between these functional identities, our dual thymidine analog labeling paradigm yielded valuable insights into the timeline of the proliferative stages of neurogenesis. Given the asynchronous nature of cell divisions observed in vivo, we opted for 3 injections spaced 9 hours apart for each thymidine analog, assuming an average S-phase length of approximately 11 hours (Fischer et al., 2014) in order to label the complete cohort of dividing cells within a 20 hour window. Our results show that cells enter and exit active neurogenic proliferation within a window of 96 hours (duration between the first CldU and IdU injections; 90% of all cells). These results are in line with published models of precursor cell proliferation (Beccari et al., 2017; Podgorny et al., 2018) and with data from longitudinal live cell imaging in vivo (Pilz et al., 2018, who reported on average 2.3 rounds of cell cycle per NSC).

Neither the distribution of Nes-GFP^+^ cells among ROS classes at baseline, nor the median cellular ROS content within individual ROS classes, nor the baseline recruitment of cells into proliferation were affected in Nox2-deficient mice. We thus propose that under standard housing conditions, the mechanisms by which a cell generates, maintains and regulates ROS is not subject to Nox2 activity, but rather that Nox2-mediated ROS fluctuations represent an independent mode of controlling cell cycle entry that can be triggered by environmental or activity-dependent cues. Acute physical activity elicited Nox2 mediated ROS fluctuations in the quiescent hiROS cells, which are necessary for an initial pro-proliferative response, reflected by the downstream redistribution of Nes-GFP^+^ cells and an increase in the loROS NPC fraction. Previous work, employing neurospheres from the SVZ and other in vivo models, has demonstrated that plasma membrane-bound Pten, a well-established tumor suppressor, acts as a downstream target of Nox2 (Le Belle et al., 2011; Hervera et al., 2018). Nox2-generated ROS leads to the oxidation and inactivation of Pten, which augments phosphatidylinositol 3,4,5-triphosphate (PI3K) production and Akt activation, which further phosphorylates a multitude of factors that govern cellular responses including cellular growth and proliferation (Luo et al., 2003). Based on our results of an absent phenotypic redistribution among IdU^+^ cells upon stimulation (Figure 1D, E) we conclude that in a naïve mouse, de novo physical activity triggers the activation of quiescent cells into an invariable scheme of neurogenic progression. Upon sustained stimulation, the acute response to physical activity may give way to an effect on maintenance of the precursor cell pool, which is in line with the observation that upon prolonged exposure to the running wheel, net neurogenesis continues to increase at a time point when proliferation levels have already receded (Kronenberg et al., 2003). In other words: a novel exercise stimulus would exert a stronger effect on a naïve population of type-1 cells than on a pool of cells that has already been primed. The neurogenic niche would acquire a memory of activation reflected in the functional states of its stem cells.

Our results further allow the extended hypothesis that, dependent on ROS-defined functional state, cells along the proposed pseudotime axis proposed by Shin et al. (Shin et al., 2015) might actually be able to shuttle between the identifiable early developmental stages and future work shall be focused on unraveling mechanisms which regulate intracellular ROS levels and lead to cellular state transition. Thus, in the adult mammalian hippocampus, which represents a late evolved and extremely specialized structure with crucial functionality (Kempermann, 2012; Treves et al., 2008; Wiskott et al., 2006), we show that a primordial mechanism for defining cellular states based on redox potential is uniquely refined to regulate homeostasis of neurogenic stages under different physiological demands and thereby regulate a highly advanced mechanism of structural plasticity.

## Supporting information

Signature_genes_marking_Nes-HGP_of DG_SVZ

Enriched_transcripts_Nes-GFP_ROS_classes

Signature_transcripts_Nse-GFP_ROS_class

Transcripts_ROS_Drops

Curated_query_gene_lists

Table_replicate_numbers

## Author contributions

Conceptualization: VSA, AER and GK; Methodology: VSA, TLW, AER, RWO, DGK; Investigation: VSA, TLW, AER, SA, GMK, TJF, SZ; Software, data curation: RWO; Formal analysis: AER, VSA; Visualization: AER, RWO; writing-original draft-VSA, AER, GK; Writing-revision-TLW, RWO, SZ, JM; Supervision: AER, JM, GK; Project administration: AER, GK; Funding acquisition: GK.

### Acknowledgments

This work was supported by the Flow Cytometry core facility of the CRTD and the Dresden Genome Center (DGC) at Technische Universität Dresden. We would like to acknowledge the support of the present and past animal keepers and technicians of the Kempermann group, especially Nicole Rund, Anne Karasinsky, Sandra Günther. DGK is a member of the Dresden International Graduate School for Biomedicine and Bioengineering (DIGS-BB) PhD program. JM is supported by the German Research Foundation (DFG) (Emmy Noether; MA 5831/1–1) and receives funding from the European Research Council (ERC) under the European Union’s Horizon 2020 research and innovation program (grant agreement no. 680042). We acknowledge the extended support by lab members and alumni of the Kempermann group. This work was funded by the VI-510 “RNA metabolism in ALS and FTD” and the AMPro consortium of the Helmholtz association.

## STAR Methods

Key Resources Table

| REAGENT or RESOURCE | SOURCE | IDENTIFIER |
| --- | --- | --- |
| **Antibodies** |  |  |
| Mouse monoclonal anti- IdU | BD Bioscience | 347580 |
| Goat polyclonal anti-doublecortin (DCX) | Santa Cruz | sc-8066 |
| Rat monoclonal anti-BrdU (CldU/BrdU) | AbD Serotec | OBT0030 |
| Rabbit polyclonal anti-Ki67 | Novocastra | NCL-Ki67p |
| Goat polyclonal anti-Sox2 | Santa Cruz | sc-17320 |
| Rabbit polyclonal anti-Tbr2 | Abcam | ab23345 |
| **Bacterial and Virus Strains** |  |  |
| **Biological Samples** |  |  |
| **Chemicals, Peptides, and Recombinant Proteins** |  |  |
| B27 supplement | Thermo Fisher Scientific | 17504044 |
| CellROX Deep Red reagent | Thermo Fisher Scientific | C10422 |
| 5-Chloro-2'deoxyuridine (CldU) | Sigma-Aldrich | C6891 |
| diaminobenzidine (DAB) | Sigma-Aldrich | D5905 |
| Dihydroethidium | Thermo Fisher Scientific | D1168 |
| Epidermal growth factor | Peptotech | AF-100-15 |
| Basic fibroblast growth factor | Peptotech | AF-100-18B |
| GlutaMAX supplement | Thermo Fisher Scientific | 35050061 |
| Heparin | MP Biomedicals | 0210193125 |
| 5-Iodo-2'-deoxyuridine (IdU) | Sigma-Aldrich | I7125 |
| Neurobasal medium | Thermo Fisher Scientific | 21103049 |
| MitoSOX | Thermo Fisher Scientific | M36008 |
| eBioscience™ Fixable Viability Dye eFluor™ 780 | Thermo Fisher Scientific | 65-0865-14 |
| Propidium Iodide | Thermo Fisher Scientific | P3566 |
| Potassium Chloride | Sigma-Aldrich | P9333 |
| **Critical Commercial Assays** |  |  |
| Neural Tissue Dissociation Kit (P) | Miltenyi Biotec | 130-092-628 |
| RNeasy Micro Kit | Qiagen | 74004 |
| VECTASTAIN ABC-HRP Kit | Vector Laboratories | PK-4000 |
| Deposited Data |  |  |
| Raw Sequence data | This paper | GSE124095 |
| Experimental Models: Cell Lines |  |  |
| Experimental Models: Organisms/Strains |  |  |
| B6.129S-Cybb<sup>tm1Din</sup>/J | The Jackson Laboratory | #002365 |
| C57BL/6JRj | Janvier |  |
| Nestin-GFP | Yamaguchi et al., 2000 |  |
| Oligonucleotides |  |  |
| Recombinant DNA |  |  |
| Software and Algorithms |  |  |
| Prism | GraphPad | V7 |
| FlowJo | FLOWJO, LLC | V10 |
| Diva | BD Biosciences | V8.01 |
| Other |  |  |

Contact for Reagent and Resource Sharing

Further information and requests for resources and reagents should be directed to and will be fulfilled by the Lead Contact Professor Gerd Kempermann.

## Experimental Model and Subject Details

### Mice

Mice were maintained on a 12-/12-h light/dark cycle with access to standard mouse chow (Sniff) and water ad libitum. Animals were aged between 6-8 weeks old at the time of the experiment, unless otherwise stated. C57BL/6JRj female mice were purchased from Janvier Labs. Nox2 knockout mice (B6.129S-Cybb^tm1Din^/J; (Pollock et al., 1995)) were initially purchased from The Jackson Laboratory and maintained as a heterozygous breeding colonies, Nestin-GFP mice (Yamaguchi et al., 2000), were obtained from Yamaguchi and colleagues and maintained as a homozygous colony, Mice were housed in groups of at least two mice in standard polycarbonate cages (Type III, Techniplast, Germany), unless otherwise stated. All experiments were conducted in accordance with the applicable European regulations and approved by the responsible authority (Landesdirektion Sachsen). A supplementary file detailing the genotype of the animals used for each experiments; number of animals per sample and sample size is available with this manuscript.

## Method Details

### Physical activity paradigm

Animals were singularly housed in either standard cages or those containing a running wheel (150 mm diameter, TSE Systems, Germany) for 2 or 4 days. The distance run was monitored and recorded.

### Thymidine labelling and tissue preparation

To label proliferating cells, animals were injected intraperitoneally with CldU (42.5 mg/kg), IdU (57.5 mg/kg) or BrdU (50 mg/kg). See individual experimental set-up paradigms for the timing of the labelling. Mice were transcardially perfused with NaCl (0.9 % w/v) and brains removed and post-fixed in 4 % paraformaldehyde (PFA) at 4 °C overnight. The next day, brains were transferred to a 30 % sucrose solution for 2–3 days. Coronal sections with a thickness of 40 µm were cut using a sliding microtome (Leica SM2010) cooled with dry ice. Sections were collected and stored as floating sections in cryoprotection solution (CPS; 25 % ethyleneglycol, 25 % glycerol in 0.1 M phosphate buffer pH 7.4) at −20°C. Every sixth section of each brain was pooled in one series for immunohistochemistry.

### Fluorescence immunohistochemistry

Two combinations of antibodies were used to phenotype the proliferating cells (Sox2/Tbr2/IdU/CldU, or Tbr2/DCX/IdU/CldU). Sections were first washed with PBS and treated with 0.9 % NaCl, before DNA denaturation was performed in 2 N HCl for 30 min at 37 °C. The sections were then thoroughly washed with PBS and blocked for 1 h in PBS supplemented with 10 % donkey serum (Jackson ImmunoResearch Laboratories Inc) and 0.1 % Triton X-100. Primary antibodies (Sox2, Tbr2, DCX, rat anti-BrdU, or mouse anti-BrdU) were diluted in PBS supplemented with 3 % donkey serum and 0.1 % Triton X-100. Incubation was performed overnight at 4 °C. After several rinses in PBS, the sections were incubated with secondary antibodies diluted in PBS supplemented with 3 % donkey serum and 0.1 % Triton X-100 at room temperature for 4 h. They were then washed in PBS, after which 4’,6-diamidino-2-phenylindole (DAPI, 1:2000) staining was performed for 10 min. After a final wash with PBS, the sections were mounted on glass slides and coverslipped with Aqua-Poly/Mount (Polysciences Europe GmbH). CldU^+^ and IdU^+^ cells in every sixth section (240 μm apart) were counted along the complete ventral dorsal extent of the DG at 40x magnification using a Zeiss Apotome microscope. Results were multiplied by 6 in order to obtain the total number of positive cells within the DG region of each brain. 100 randomly-selected CldU^+^ and IdU^+^ cells per DG were photographed using the apotome function and phenotyped for the co-expression of Sox2 and Tbr2; and Tbr2 and DCX.

### DG and SVZ dissection and dissociation

Mice were killed, their brains immediately removed, and the dentate gyrus (DG) and SVZ microdissected (Hagihara et al., 2009; Walker and Kempermann, 2014). The tissue was enzymatically digested using the Neural Tissue Dissociation Kit (Miltenyi) according to the manufacturer’s instructions. Following a final wash in Hank’s balanced salt solution (HBSS) (PAA; GE Healthcare) the pellet was resuspended in 1 ml of growth medium or HBSS, based on the succeeding procedure, and filtered through a 40 μm cell sieve (Falcon; BD Biosciences).

### Flow cytometry

Dissociated SVZ or DG cells were analysed using a FACS Aria III cell sorter (BD Biosciences) using a 85 μm nozzle. Cells were first gated based on the forward and side scatter (FSC and SSC) to identify the main cell population. Dead cells were excluded using the live/dead dye (vitality dye eFluor™ 780 in conjugation with DHE and Mitosox, PI in conjugation with CellROX DeepRed) and doublets were excluded based on the height and width of the cells. Cells were then gated based on their ROS content. For intracellular ROS measurements, either CellROX DeepRed Reagent (5 μM, Thermo Fisher Scientific, for the neurosphere assays) or dihydroethidium (DHE; 5 μM, Thermo Fisher Scientific, for all other ROS analyses) was added to each well and incubated for 30 min at 37 °C. For mitochondrial ROS measurements MitoSOX Red Mitochondrial Superoxide indicator (5 μM, Thermo Fisher Scientific) was added and incubated for 10 mins. The cells were transferred to a falcon tube and washed with 5ml of PBS. Following centrifugation, the supernatant was removed, the cells were resuspended in 200 µl PBS and transferred to a FACS tube for sorting. Cell populations were sorted into either medium (for neurosphere experiments), RLT buffer (for RNA isolation) or PBS (western blot analyses).

### Gating for ROS classes

The cells of the DG and SVZ yielded very stereotypical ROS profiles, which were manually clustered, based on the density of clustering cells, into 4 non-overlapping classes using DIVA (BD Biosciences) for sorting the populations or during analysis using FlowJo (V10). The gates obtained from standard housed DG were used as the “master gates” which were applied to runner DG cells’ profiles or to the SVZ cells.

### Neurosphere culture

For neurosphere culture, cells were resuspended in 10 ml neurosphere growth medium consisting of Neurobasal medium (Gibco, Life Technologies), supplemented with 2 % B27 (Invitrogen), 1X GlutaMAX (Life Technologies) and 50 units/ml penicillin/ streptomycin (Life Technologies). The following growth factors were also included: 20 ng/ml EGF, 20 ng/ml FGF-2 and 20 ng/ml Heparin. Cells were plated into 96 well plates and incubated at 37 °C with 5 % CO2 and the resulting neurospheres counted using an inverted light microscope after either 7 days for SVZ cultures or 14 days for DG cultures. We used 15 mM KCl or L-(-)-noradrenaline (+)-bitartrate salt monohydrate (norepinephrine; 10 μM) as a stimulant to trigger latent cells to form neurospheres in vitro.

#### Next Generation sequencing

### RNA extraction and sequencing

Two separate RNA sequencing experiments were performed in this study. In the first experiment, tissue was microdissected from the SVZ and the DG of the same animal following the protocol of Walker et al. (2014). RNA was extracted using the RNeasy micro kit (Qiagen). In the second experiment, Nes-GFP positive cells from the DG were gated into ROS classes, based on their intracellular ROS content. 400 GFP positive cells from each ROS class were FACsorted into a PCR tube containing 8.5 *μ*l of a hypotonic reaction buffer and RNA was prepared using the SMARTer Ultra Low RNA HV Kit (Takara Bio) according to the manufacturer’s protocol. For both experiments, cDNA of polyadenylated mRNA was synthesized from RNA of the lysed cells using SmartScribe reverse transcriptase, a universally tailed poly-dT primer and a template switching oligonucleotide (Takara Bio). This was followed by 12 cycles of amplification of the purified cDNA with the Advantage 2 DNA Polymerase (Takara Bio). After ultrasonic shearing of the amplified cDNA (Covaris S2), samples were subjected to standard Illumina fragment library preparation using the NEBnext Ultra DNA library preparation chemistry (New England Biolabs). In brief, after physical fragmentation by ultrasonication (Covaris LE 220) cDNA fragments were end-repaired, A-tailed and ligated to indexed Illumina Truseq adapters. Resulting libraries were PCR-amplified for 15 cycles using universal primers, purified using XP beads (Beckman Coulter) and then quantified with the Fragment Analyzer (Advanced Analytics/Agilent). Final libraries were equimolarly pooled and subjected to 75 bp single-end sequencing on the Illumina HiSeq2500 platform, providing ~35 (24–60) million reads per sample. Reads were mapped to the latest mouse genome build (mm10) using the STAR algorithm (Dobin et al., 2013) and counts per ENSEMBL gene model prepared using the *RSubread* package (Liao Y et al., 2013) or R/BioConductor.

### Quality control and differential expression

RNASeq counts were filtered to have at least 1 count per million reads (CPM) in a minimum of 75 % of the samples from at least one cell population. CPM were calculated using the function *cpm* from the *edgeR* package in R/BioConductor (Robinson et al., 2010). Samples were clustered by plotting the first two principal components and by unsupervised hierarchical clustering. One sample from each of the two experiments showed reduced sequencing depth and complexity and did not cluster with replicates. In both cases, these samples came from preparations with very low input RNA so we decided to remove them from further analyses. A filter for differential expression was then performed using the *edgeR* functions *lmFit* and *topTags* and only significantly (adjusted *p* < .05) differentially expressed transcripts were used for enrichment analyses. Transcripts were classified as ‘enriched’ in an experimental group if the mean expression in that group was significantly above the average expression over all groups. Enriched genes were further filtered by hierarchical clustering into two clusters based on inter-group *t*-statistics and those where only one group clustered separately were termed ‘signature’ genes (for the DG/SVZ comparison, these are obviously equivalent to the ‘enriched’ gene set).

### Functional enrichment and expression profiles

Enrichment for Gene Ontology terms was performed using the R package *topGO* (Alexa et al., 2018) with the DG/SVZ ‘enriched’ gene lists as query sets and all genes passing the CPM filter (see above) as background. Expression profiles of curated gene lists were calculated by identifying all genes in the current data that corresponded to the genes in the gene list of interest and calculating the first principal component to collapse their expression into an ‘eigengene’. The eigengene values were then used for plotting. A similar approach was used to reanalyse the Shin et al. dataset where the genes corresponding to the ‘signature’ genes for each ROS class were identified in the single-cell dataset and the first principal component of these used to create an eigengene. Smooth spline interpolation of the eigengene was then performed to produce the plotted values.

### Ki67 immunohistochemistry and quantification of in vivo proliferation

Briefly, brain sections stored in CPS were transferred into PBS and washed. Endogenous peroxidase activity was blocked by adding 0.6 % hydrogen peroxide (H_2_O_2_; Merck Millipore) for 30 min and sections were then rinsed with 0.9 % NaCl. Protein-binding sites were blocked with a blocking solution (10 % donkey serum, 0.2 % Triton X-100 in PBS) for 1 h. Ki67 staining was performed with the Ki67 primary (rabbit anti-Ki67, 1:500; Novocastra) and donkey anti-rabbit-biotin secondary antibodies (1:1000; Jackson Immunoresearch Laboratories). Detection was performed using the Vectastain ABC-Elite reagent (Vector Laboratories) with diaminobenzidine (Sigma-Aldrich) and 0.04 % NiCl as the chromogen. Sections were mounted onto gelatin-coated glass slides, dried, cleared with Neoclear (Merck) and coverslipped using Neo-mount (Merck). Every sixth section (240 μm apart) was counted in the complete ventral dorsal extent of the dentate gyrus, at 40x magnification using a standard brightfield microscope. Results were multiplied by 6 in order to obtain the total number of positive cells within the dentate gyrus region of each brain.

### Quantification and Statistical Analysis

Data analysis (with the exception of the next-generation sequencing data) was performed using Prism software (Version 7, GraphPad Software, Inc). Flow cytometry data was analysed using the FlowJo software. Results were expressed as mean ± standard error of the mean (SEM). Statistical significance was determined using a Student’s t-test when the experiment contained two groups, or an ANOVA when comparing more than two groups. Dunnett, Tukey’s and Sidak post hoc tests were performed and mentioned in the text wherever applicable. The level of conventional statistical significance was set at *p* < .05 and displayed visually as * *p* <0.05, ** *p* < 0.01, *** *p* < 0.005 and **** *p* < 0.001. The number of mice or repeat experiments performed per group is stated in a separate supplementary file.

### Data and Software Availability

The raw sequence data for both the DG/SVZ and ROS experiments are deposited in GEO under the SuperSeries accession number GSE124095.

**Supplementary File 1:** Signature transcripts of Nes-GFP of DG and SVZ with functional enrichment results from Gene Ontology and Reactome.

**Supplementary File 2:** Enriched transcripts of Nes-GFP of different ROS classes in the DG with functional enrichment results from Gene Ontology and Reactome.

**Supplementary File 3:** Signature transcripts of Nes-GFP of different ROS classes in the DG with functional enrichment results from Gene Ontology and Reactome.

**Supplementary File 4:** Transcripts significantly changing at each ROS ‘drop’ with functional enrichment results from Gene Ontology and Reactome.

**Supplementary File 5:** Curated gene lists used for identification of cellular state.

**Supplementary File 6:** Table of replicate numbers for all experiments.

## Supplementary Figures with legends

**Figure S1:**
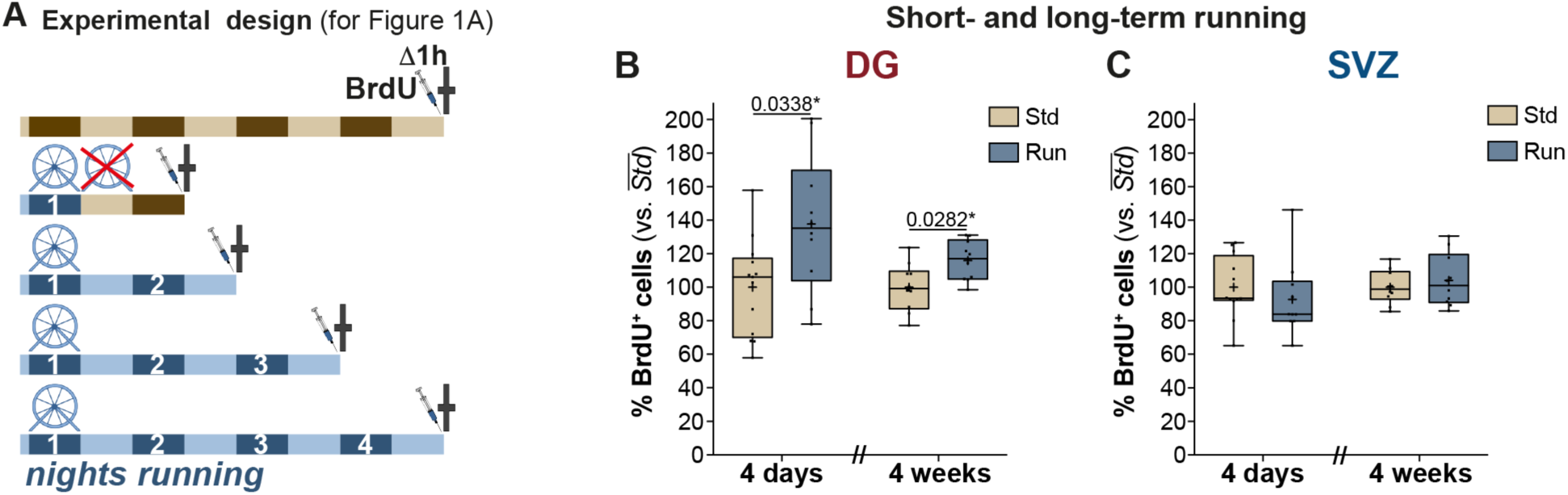
**(A)** Experiment outline for the paradigm analysing the number of proliferating cells, marked by BrdU, post varying bouts of physical activity (result shown in Fig 1A). Relative comparison of BrdU positive cells post a 4-day stimulus (short-term running) and a 4-week stimulus (long-term running) in the DG **(B)** and the SVZ **(C)**. Note that while both paradigms significantly increase the pool of proliferating cells in the DG, a similar effect could not be observed in the SVZ. Unpaired t-tests with Bonferroni corrections. All data represent mean ± SEM. * *p* < 0.05.

**Figure S2:**
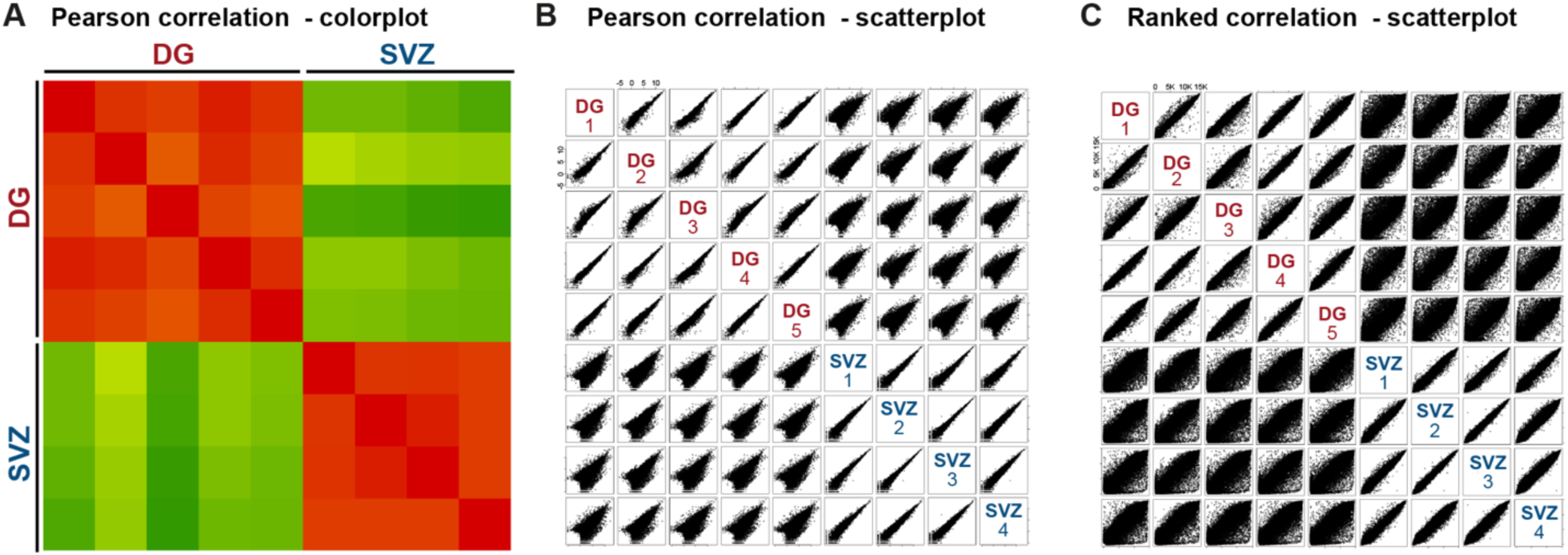
Correlation analysis between the expression profiles of complete pool of the Nes-GFP^+^ of the DG and SVZ. (A) Pearson correlation colour plot between the samples collected from separate experiments; (B) scatterplot of the Pearson correlation between the samples; (C) scatterplot of the Ranked correlation between samples.

**Figure S3:**
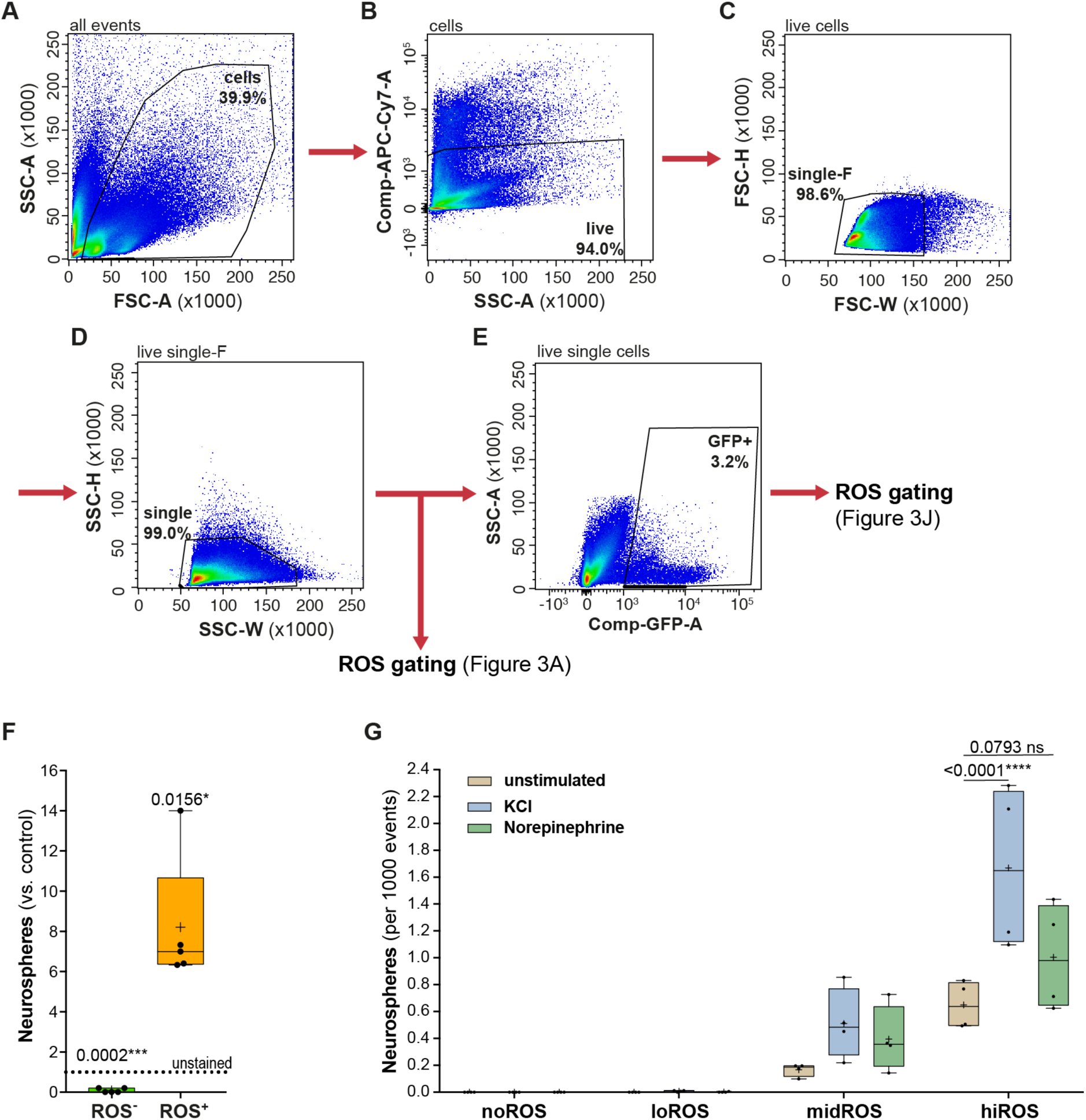
Pipeline for cytometric analyses and gating strategies (A-E). All events were gated for forward scatter (FSC-A, marks the size of events) and sideward scatter (SSC-A, marks the granularity) to exclude debris (A); These events were gated to identify putative live cells by the exclusion of vitality dye e780 (APC-Cy7 filter) and marked as live cells (similar strategy used for PI as a vitality dye) (B); The live cells were gated on size (C) and granularity (D) to identify singlets. The singlets were gated for ROS content, as shown in Figure 3A and independently gated for cells expressing GFP (E); (F) All the isolated cells of DG were gated into either ROS+ class or ROS-class and assayed for neurosphere formation. Neurosphere formation is restricted to the higher ROS class, and significantly enriched compared to unstained sorted events. One-sample t-test with Bonferroni correction; (G) Plot showing the neurosphere formation from each of the ROS class with or without stimulation with KCl (blue) and Norepinephrine (NE, green). Irrespective of the stimulus, neurosphere formation was restricted to mid and high ROS classes. Stimulation with with either KCl or NE does not significantly increase neurosphere formation in the midROS class. However, stimulation with KCl strongly augments more neurosphere formation in the hiROS class (p < 0.0001) and stimulation with NE increases neurosphere formation to a lesser extent (p = 0.0793). Two-way Anova with Dunnett post-hoc correction. All data represent mean ± SEM. * *p* < 0.05, **** *p* < 0.0001.

**Figure S4:**
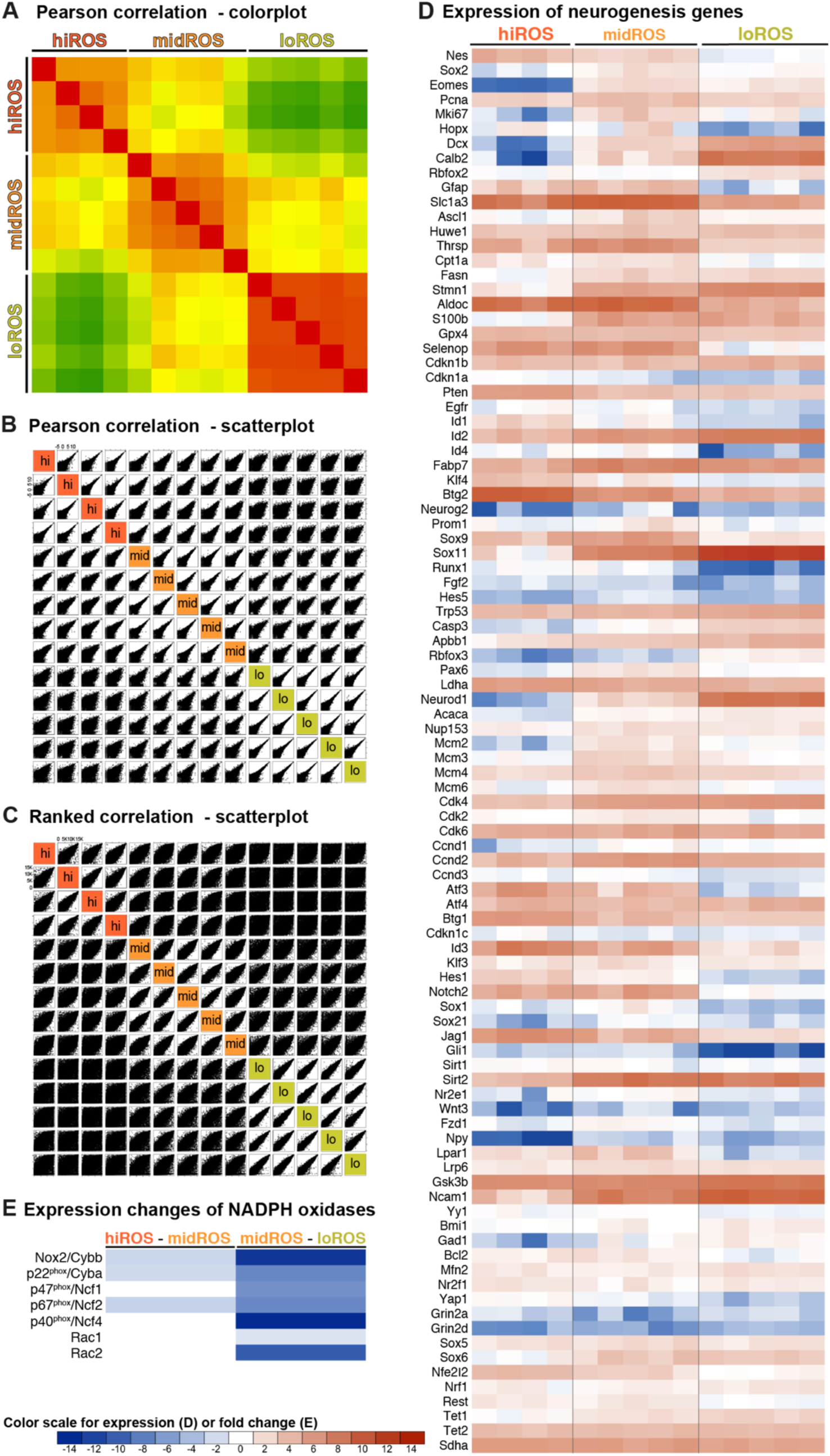
Correlation analysis between the expression profiles of the Nes-GFP^+^ of the different ROS classes within the DG (hiROS class indicated in red, midROS in orange and loROS in green) (A) Pearson correlation colour plot between the samples collected from separate experiments; (B) scatterplot of the Pearson correlation between the samples; (C) scatterplot of the Ranked correlation between samples; (D) heatmap showing the expression levels and the fold changes of curated transcripts, with a known role in neurogenic maintenance and progression and cell cycle regulation, in the Nes-GFP^+^ of the three ROS classes (see supplementary file 3 for the raw values); (E) heatmap showing the changes in the expression of Nox complex genes at the transition of cellular ROS content from high to mid (left) and from mid to low (right). Colour scheme indicates significance of change (white indicates no significant change) and the degree of fold change.

**Figure S5:**
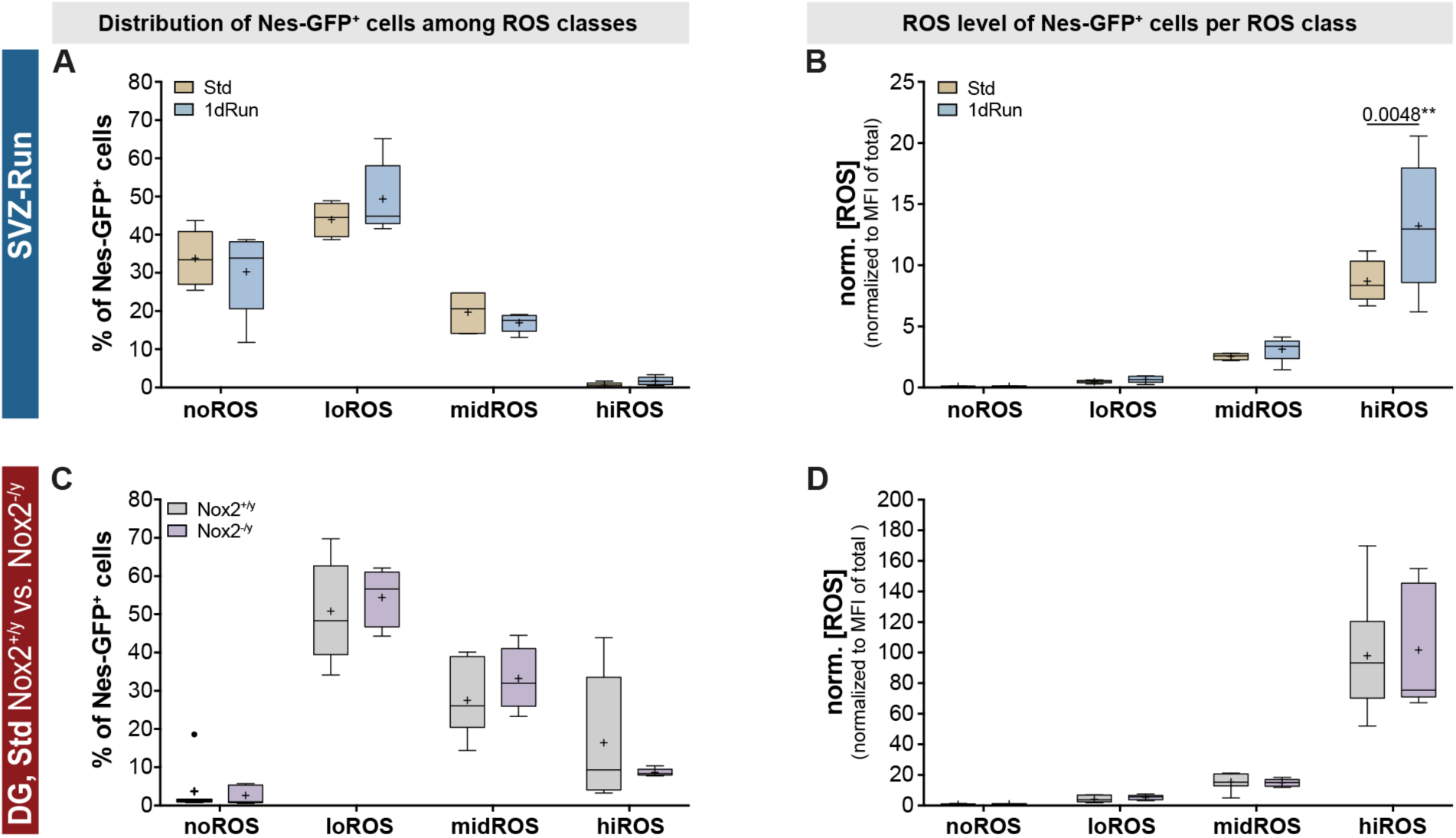
Plots showing the changes in the distribution of Nes-GFP^+^ cells within the SVZ into the different ROS classes (A) and the changes in their median intracellular ROS content (B) post 1-day running paradigm. Note that although a ROS surge, similar to the surge observed in hiROS Nes-GFP^+^ cells of the DG, could be observed in the SVZ, the proportion of this population is significantly smaller in SVZ (<1%). Two-way ANOVA with Sidak post-hoc test. Plots showing the distribution of Nes-GFP^+^ cells within the DG into the different ROS classes of Nes-GFP::Nox2^-/y^ mice (Nox2 mutants) and littermate Nes-GFP::Nox2^+/y^ controls (wildtype controls). Two-way ANOVA with Sidak post-hoc test. All data represent mean ± SEM. * *p* < 0.05, ** *p* < 0.01, *** *p* < 0.001, **** *p* < 0.0001.

